# Optimizing Bioprocessing Efficiency with OptFed: Dynamic Nonlinear Modeling Improves Product-to-Biomass Yield

**DOI:** 10.1101/2024.07.31.605953

**Authors:** Guido Schlögel, Rüdiger Lück, Stefan Kittler, Oliver Spadiut, Julian Kopp, Jürgen Zanghellini, Mathias Gotsmy

**Affiliations:** Department of Analytical Chemistry, University Vienna, Währinger Straße, 1090 Vienna, Austria, EU; Doctorate School of Chemistry, University of Vienna, Währinger Straße, 1090 Vienna, Austria, EU; Integrated Bioprocess Development, Technical University Vienna, Getreidemarkt 9, 1060 Vienna, Austria, EU; Austrian Centre of Industrial Biotechnology, Krenngasse 37, 8010 Graz, Austria, EU

**Keywords:** fed-batch, bioproduction, non-linear optimization, process design

## Abstract

Biotechnological production of recombinant molecules relies heavily on fed-batch processes. However, as the cells’ growth, substrate uptake, and production kinetics are often unclear, the fed-batches are frequently operated under sub-optimal conditions. Process design is based on simple feed profiles (e.g., constant or exponential), operator experience, and basic statistical tools (e.g., response surface methodology), which are unable to harvest the full potential of production.

To address this challenge, we propose a general modeling framework, OptFed, which utilizes experimental data from non-optimal fed-batch processes to predict an optimal one. In detail, we assume that cell-specific rates depend on several state variables and their derivatives. Using measurements of bioreactor volume, biomass, and product, we fit the kinetic constants of ordinary differential equations. A regression model avoids overfitting by reducing the number of parameters. Thereafter, OptFed predicts optimal process conditions by solving an optimal control problem using orthogonal collocation and nonlinear programming.

In a case study, we apply OptFed to a recombinant protein L fed-batch production process. We determine optimal controls for feed rate and reactor temperature to maximize the product-to-biomass yield and successfully validate our predictions experimentally. Notably, our framework outperforms RSM in both simulation and experiments, capturing an optimum previously missed. We improve the experimental product-to-biomass ratio by 19 % and showcase OptFed’s potential for enhancing process optimization in biotechnology.

## 1. Introduction

Biotechnological production processes are the backbone of numerous industries, from pharmaceuticals to biofuels. Many of these processes are operated as a fedbatch, adding substrate and nutrients continuously when the initial batch medium is depleted [1]. This method allows for control of key parameters, such as nutrient concentration, and is fundamental to achieving high yields and product quality. Consequently, the optimization of such processes becomes a critical objective.

Optimizing fed-batch processes is challenging due to the complexity of cellular mechanisms, which are difficult to measure directly and can vary significantly based on factors such as product type, microorganism, induction mechanism, and product location. As a result, optimization often involves a trial-and-error approach [2]. In this context, theoretical and mathematical modeling offers a powerful and complementary alternative. By leveraging simulations and model-based design of experiments [3, 4], we can reduce our dependence on costly and time-consuming trials, and enhance our ability to predict and optimize process performance in a more controlled and efficient manner.

In general, there are two distinct paths for optimizing biotechnological processes: statistical design of experiments, such as response surface methodology (RSM) [5], and the model-based approach [3, 4, 6, 7]. Statistical methods, including RSM, offer straightforward and accessible means of optimization. However, they do not leverage biological knowledge, which could enhance the optimization process [8], and are limited to optimizing a predefined set of discrete variables [9]. In contrast, the model-based approach begins by understanding the underlying process, representing it accurately without relying on mathematical assumptions like the quadratic dependencies used in purely statistical methods [10].

For the model-based approach, empirical models are typically employed, often formulated as ordinary differential equations (ODEs). For instance, Monod’s widely used model for population growth [11] is one such example. Product creation in these models is often linked to growth rate [4, 12, 13] or feed rate [14]. These models provide a simple yet effective means of understanding and predicting bioprocess behavior, particularly when dealing with the often limited and noisy data typically obtained from bioprocesses.

However, simplicity comes at a cost. The inherent simplifications in these empirical models constrain their applicability, limiting their ability to describe complex processes comprehensively. These straightforward models often fail to account for some phenomena observed under constant process conditions on a cellular level. This includes production deterioration over time due to metabolic adaptation, or product inhibition [15]. Despite their limitations, these models are frequently used because formulating more complex models is hindered by sparse data, biological variation, and the difficulties in selecting the best available model equations [16]. Different metrics can be used for model selection, e.g., AIC and AIC_C_ [17], LASSO [18], and methods based on cross-validation [19, 20]. Regardless of the used metric, the number of possible models can get very large (e.g., over 2^18^ ≈ 2.6 × 10^5^ possible models with 18 parameters) which requires defined search strategies (e.g., HIPPO) [21, 22]. Moreover, decisions in the modeling process can introduce biases that influence the outcome [23, 24], and using more complex models with small datasets may lead to overfitting [25].

The modeling process does not exist in isolation; its primary goal is to improve the efficiency of the process (e.g., by maximizing titer/biomass or minimizing operational costs). This objective often involves defining specific targets for each process, frequently utilizing the TRY metric (titer, rate, yield) [26]. The choice of optimization algorithms varies depending on the complexity of the model at hand, which is influenced by the available data set and its quality. While straightforward maximization algorithms suffice for discrete variables [27], optimizing continuous solutions, such as feed and temperature functions, poses mathematical challenges, notably due to the theoretically infinite number of control variables [28]. To address these challenges, various mathematical methods have been developed. For simpler models, Euler-Lagrange equation-based approaches can be employed [4]. In contrast, more intricate models, as encountered in our work, require the application of optimal control theory [29].

The realm of optimal control problems has been extensively explored, resulting in a multitude of solution methods [30]. Analytical methods, such as those grounded in Pontryagin’s maximum principle [29], are well-suited for relatively straightforward problems but may not be practical for complex real-life scenarios. In most cases, numerical solvers become essential. The two primary categories of solvers are direct methods and those based on dynamic programming. Direct methods tackle the optimization and differential equations simultaneously, transforming the problem into a set of nonlinear differential equations [31]. In contrast, dynamic programming [32, 33] relies on the principle of optimality within Hamilton-Jacobi-Bellman frameworks, iteratively solving the problem. While dynamic programming holds the promise of identifying global optima, it tends to be slower than direct methods and is often infeasible for high-dimensional problems. As a result, direct methods find more frequent applications, particularly in engineering contexts, where a wealth of software packages is available for implementation [34–37].

In this study, we present OptFed, a comprehensive framework using an ODE model to describe bioprocesses. The framework is divided into three stages: define, fit, and optimize. First, we define a general and flexible form of the ODE model. Next, its kinetic parameters are fitted to training data and the model size is reduced to avoid overfitting. To do this, we developed a heuristic algorithm that starts with the general model and removes terms and parameters that do not significantly improve the fit. In the third stage, based on the reduced model, we leverage optimal control theory to identify optimal values for control variables. In a case study, we apply OptFed to protein L production to maximize the product-to-biomass yield. Optimal values for the temperature and substrate feed control are predicted. A comparison to RSM highlights the improvements of OptFed to typically used statistical methods. Moreover, experimental validation results in a 19 % improved product-to biomass yield.

## 2. Methods

### 2.1. Modeling Framework

Our modeling framework comprises three key components:

I. define,
II. fit, and
III. optimize.

In the first stage, we establish a general process model capable of representing a wide range of biotechnological (fed-)batch production processes. In the second stage, the general model is fitted to specific process data, the kinetic parameters are estimated and insignificant terms are removed. In the third stage, the fitted process model is used to optimize control variables, ultimately maximizing a freely selectable objective function. A graphical overview of the modeling framework is given in Figure 1.

**Figure 1:**
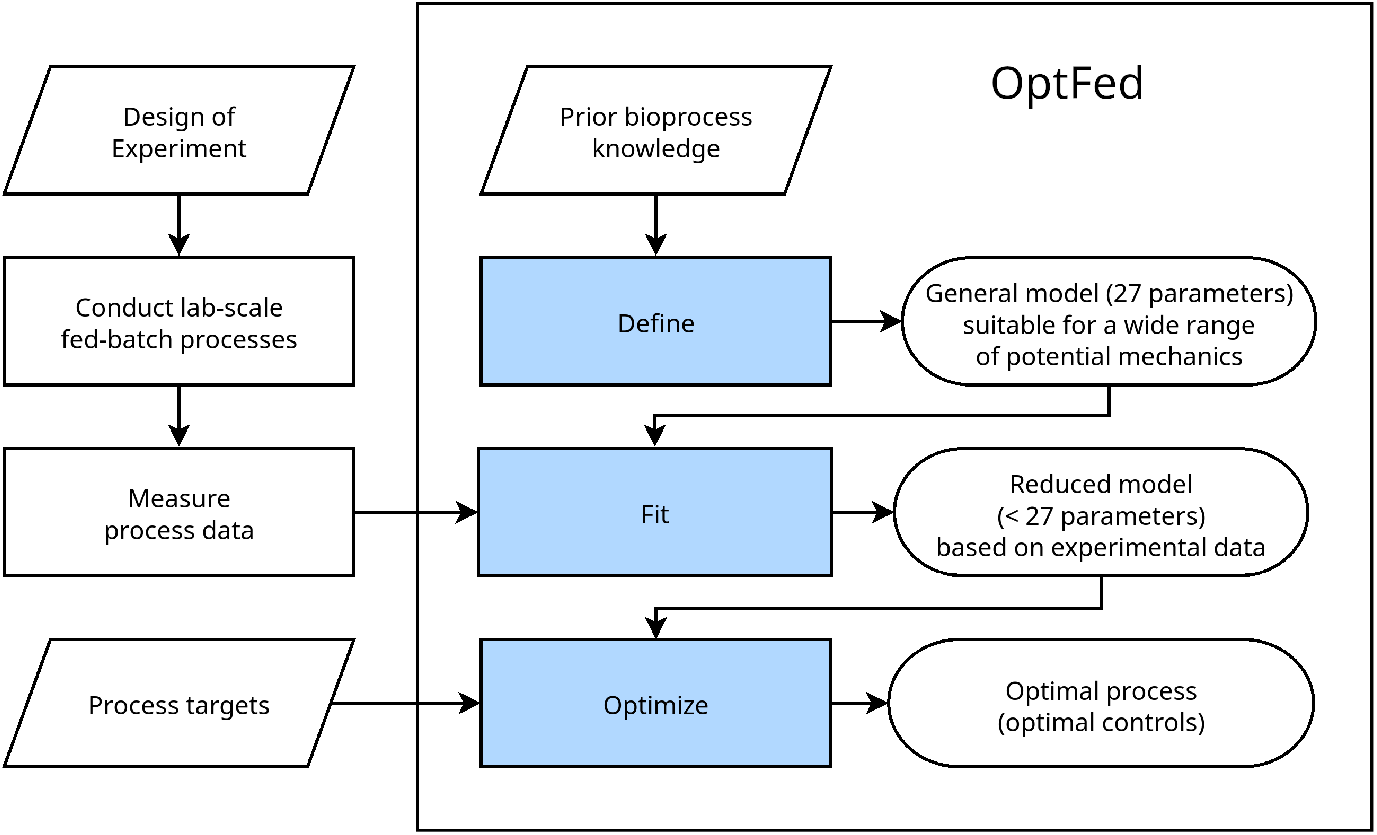
Flow chart for OptFed. In this study, we focus on the stages highlighted in blue. Initially, we set up a general model including different inhibition effects (Define). This model can describe different processes and uses many model parameters. In the second stage, the model is simplified to avoid overfitting and parameters are estimated (Fit). In the third stage, optimal process control variables are predicted (Optimize).

A list of all parameters and their respective symbols and units is given in Supplementary Table S1.

#### 2.1.1. Stage I: Define

We consider the production of recombinant proteins by *E. coli* in a fed-batch process, described by the following standard system of differential equations [1]:

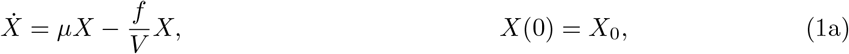

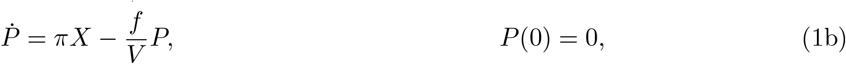

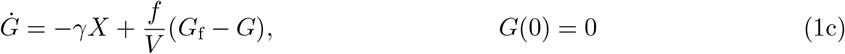

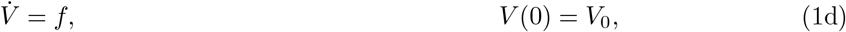

where *G, G*_f_, *P*, and *X* represent substrate concentrations in the reactor, substrate concentration in the feed, product, and total biomass, respectively, and *V* the current reactor volume. *f, γ, µ*, and *π* denote the feeding rate, substrate uptake rate per biomass, biomass growth rate, and specific product formation rate, respectively.

As the product is part of the biomass, we additionally define the metabolic active residual biomass (given as dry weight)

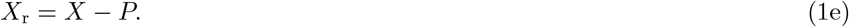

This defines the uptake per active residual biomass (*γ*^*°*^) as

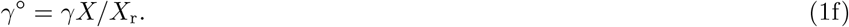

To connect the substrate uptake behavior with cellular growth and production, we assume that the total uptake can be divided into three additive components

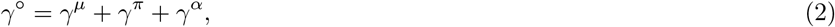

where *γ*^*µ*^, *γ*^*π*^, and *γ*^*α*^ denote the specific substrate uptake supporting growth, product formation, and cellular maintenance, respectively. Here, maintenance summarizes all cellular processes not linked to growth or production.

Using biomass-to-substrate yield 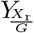 and product-to-substrate 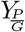, we can connect glucose consumption to product formation and growth as follows:

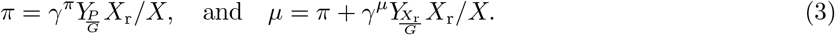

The factor *X*_r_*/X* accounts for the fact that only the metabolically active residual biomass *X*_r_ contributes to additional product formation and growth.

We assume that total substrate uptake as well as substrate uptake for product formation follow a non-competitively inhibited Michaelis–Menten process [14, 38]

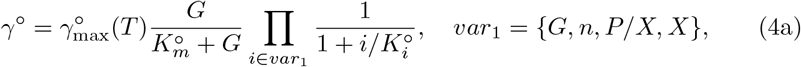

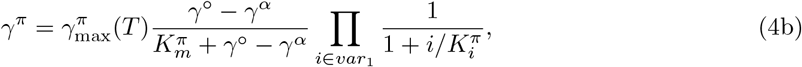

while substrate demand for maintenance is given by [39]

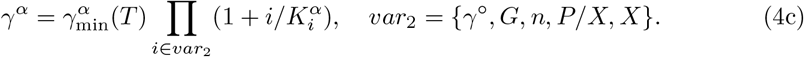

With these assumptions, *γ*^*µ*^ follows according to (2) to

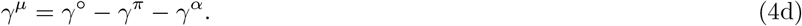

Here, 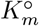, and 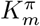 are Michaelis-Menten constants, while 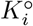 and 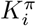 are inhibition constants, and 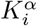 are activation constants, *n* is the number of generations (*n* = log_2_(*XV/*(*X*_0_*V*_0_))). Finally, the minimum uptake rate 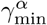 and the maximum uptake rates 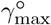 and 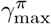 are assumed to be temperature (*T*) dependent (excluding enzyme denaturation) [40]

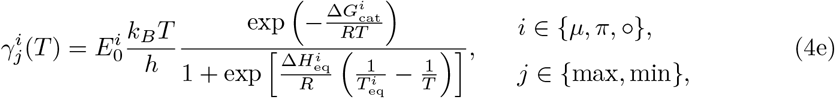

where 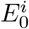 is a (hypothetical) enzyme concentration, 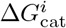, the activation energy, 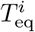 is the temperature where half of the enzymes are in an active state (the other half is inactivated by the high temperature), and 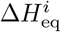 determines how abruptly the reaction rate declines with rising temperatures. The superscripts °, *α*, and *π* differentiate variables for substrate uptake, maintenance, and production rate, respectively, differentiating the constants for *γ*^*°*^, *γ*^*α*^ and *γ*^*π*^.

Our model, defined in equations (1) and (4), contains 29 free parameters. The substrate yields 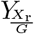, and 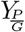 can be derived from genome-scale metabolic models [41], while the remaining 27 parameters need to be fitted from training data.

##### Bioreactor volume estimation

Generally, change in the bioreactor volume is affected by five factors, the substrate feed, the base feed (for pH control), experimental sampling, the antifoam feed, and gaseous exchanges,

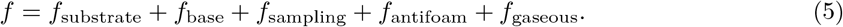

In OptFed, we explicitly model the first three of them. While the substrate feed is kept variable (for optimization), the base feed is calculated as

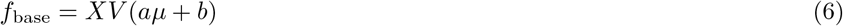

where the parameters *a* and *b* were fitted to the training data. Additionally, we accounted for volume change through experimental sampling in OptFed. For simplicity, and due to the fact that antifoam, feed, and gaseous exchanges (i.e., evaporation, O_2_ uptake, and CO_2_ excretion) are either minor or antagonistic contributors to volume change, we did not consider them in OptFed. More details on volume calculation are given in Supplementary Methods S1.1.

##### Substrate feed types

Although OptFed is not restricted to a certain type of substrate feed, it can make sense to enforce them either for comparison to the training data or for simplification of experimental implementation. Here we present two feed types that we refer to throughout the manuscript.

An exponential feed is generally very popular as the cells’ internal metabolic fluxes are (approximately) constant. It is calculated as

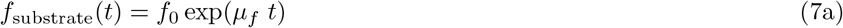

where *µ*_*f*_ of the unit h^−1^ is the defining parameter. *f*_0_ is usually derived from the properties of the process at hand.

Additionally, we also use a linear feed rate, calculated as

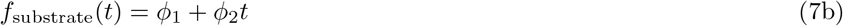

where *ϕ*_1_ and *ϕ*_2_ may be varied.

#### 2.1.2 Stage II: Fit

In this step, we use training data to select a model of suitable size and fit its kinetic parameters.

##### Training Data

OptFed requires process data for different feed rates and temperatures. Data from a central composite design [10] commonly used for RSM proofed useful (Section 2.4).

##### Experimental Data Interpolation and Rate Calculation

The differential method [42, 43] is used to estimate uptake, growth, and production rates by fitting the concentration data and differentiating the fits. We fit splines, which are continuous in both their values and first derivatives, using SciPy’s `Univariate Spline` function onto the experimental data points for control and state variables. By inserting these splines into equations (1) and (2), we calculate the experimental values for 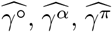 and 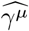 (Supplementary Methods S1.1). The hat notation indicates that these variables are derived from the experimental training data and are used to estimate the unknown parameters in (4).

##### Model Parameter Estimation

Model selection is based on a heuristic algorithm. It is inspired by ANOVA [45] and uses cross-validation [19, 20] for hyperparameter selection. By approximating uptake, growth, and production rates separately, we deal with three smaller models instead of one large, simplifying the estimation. Parameter identification is performed using differential evolution [46] with SciPy’s `differential_evolution` function [44], utilizing the previously calculated 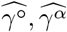 and 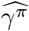 Each of the three rates is fitted separately.

We assume that the effects of each variable in (4) are independent, meaning each effect is a separate term in the equation. Each model term, containing one influencing variable and one or more parameters, can be removed (if they are deemed insignificant) and the model remains valid. In case there is no temperature effect, Eqn. (4e) simplifies to

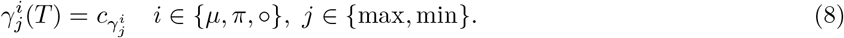

Each model term of Eqn. (4) is tested. If it does not significantly improve the model fit, it is removed according to the following algorithm:

1. **Initial Fit**: Fit the model with all currently considered parameters by minimizing the sum of quadratic errors over all processes and measurement points. Calculate the error residuals and total variance for the measurement points (Bounds used in the error minimization are shown in Supplementary Table S2).
2. **Leave-One-Out Fit**: Repeat the fitting process for models, each missing one parameter.
3. ***F* -Test**: Use an *F* -test to determine if the reduction in variance is significant (i.e., *p < α*) and calculate the difference in variance with and without the parameter.
4. **Remove Insignificant Parameters**: Remove the parameter with the highest *p*-value in the *F* -test.
5. **Iterate**: Repeat 1 to 4 until only significant terms remain.

Steps 1-5 are performed separately for each fitted rate (*γ*^*°*^, *γ*^*α*^, and *γ*^*π*^) and for 13 levels of *α*. The significance level depends on the training data (such as the number of processes and measurement points) and is computed through cross-validation. The significance level that results in the lowest error for the target variable 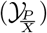 is selected (Supplemental Methods S1.2).

As for each iteration of our algorithm (Step 5), a term is removed from the model equation, the maximum amount of iterations is defined by the maximum amount of removable terms per rate equation plus one (i.e., seven).

To compare our heuristic approach to model selection with the corrected Akaike information criterion (AIC_C_) [17], we fit all possible parameter combinations for *γ*^*°*^, *γ*^*α*^, and *γ*^*π*^. Next, we calculate the AIC_C_ [Eqn. (14)] for each of the resulting models and selected the one with the lowest value.

#### 2.1.3 Stage III: Optimize

To optimize

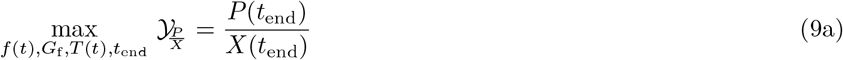

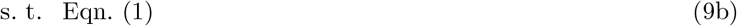

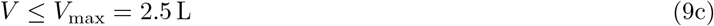

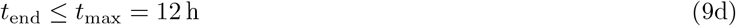

we use `ipopt` 3.14.10 [47], which is a general nonlinear programming solver. For discretization and numerical differentiation [Eqn. (1)], we implemented an orthogonal collocation and optimization algorithm in `casadi` 3.3.5 [34] using Python. All required code to reproduce our analysis is available at https://github.com/gschloegel/OptFed.

To solve the differential equations in (1), we first scale the time coordinate by setting *t* = *t*_end_*τ*, using the process’ time *t*_end_ as a control variable. We then apply orthogonal collocation on 100 finite elements [48]. Specifically, we use Gauss–Legendre polynomials of degree one with collocation points at 0.5. For the substrate, due to system stiffness, we use Gauss–Radau collocation points at 1. Locally, we solve the differential equation using the backward Euler method. The controls (feed and temperature) are linear on each of the 100 intervals, allowing the optimization method to find optima with temperature gradients and unconventional feeding strategies.

To avoid rapid variations in the control variables *u* = (*f, G*_f_, *T, t*_end_), specifically in the feed and temperature profile *f* (*t*_end_*τ*), and *T* (*t*_end_*τ*), we add a regularization term to the objective function in (9a). This modified objective reads

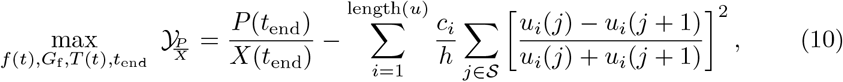

where *c*_*i*_ is the penalty factor for each control variable *u*_*i*_, *h* is the length of the finite elements, and S is the set of all sampling times.

In addition to the general optimization problem in (9), we defined a simplified version where all substrate is immediately used and no substrate accumulates, i.e., 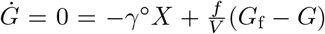 This mirrors the standard assumption in fed-batch processes and computationally avoids issues posed by stiff differential equations.

### 2.2. Response Surface Methodology

To benchmark OptFed, we evaluate its performance against response surface methodology (RSM) in process optimization. We calculated RSM using the `rsm` package [49] for R [50].

RSM requires that the control variables remain constant throughout the process and uses the process target metric (such as production concentration, yield, or productivity) to fit the model. Thus the RSM model is represented as:

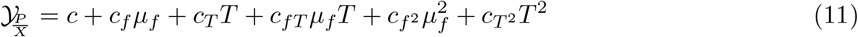

where *c, c*_*f*_, *c*_*T*_, 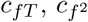, and 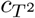 are fitted from the data minimizing the sum of quadratic errors. *µ*_*f*_ is the growth rate of the exponential feed (in 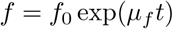).

The same model is calculated with the target variables *P* and *X*.

### 2.3. Model Comparison and Validation

To evaluate model fit [Eqn. (1)], individual (state) variables (e.g. *P, P/X, X*) or specific rates (e.g., *γ*^*°*^, *γ*^*α*^, and *γ*^*π*^), we use the coefficient of determination *R*^2^ and its adjusted version 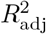, which are defined as:

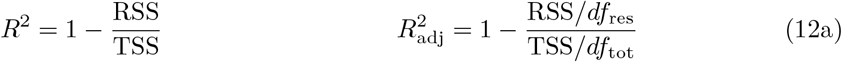

with the residual and total sum of squares (RSS and TSS)

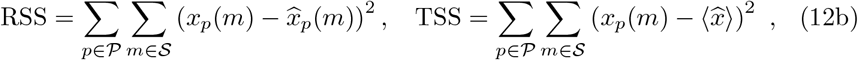

respectively. Here, *x*_*p*_ (*m*) and 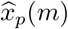 represent the predicted and measured values respectively, ⟨*x*⟩ represents the average of the observed values, P is the set of all processes, and S is the set of all sampling times. *df*_res_ and *df*_tot_ are the residual and total degrees of freedom, given by *df*_res_ = #*p* − #*v* − 1 and *df*_tot_ = #*p* − 1, respectively, where #*p* = |P|+|S| represents the number of points and #*v* represents the number of variables.

Finally, we measure relative errors with respect to the mean of all data points:

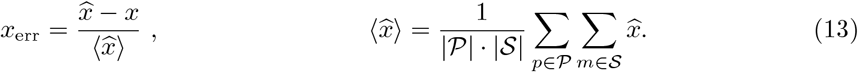

Different models are compared using *R*^2^ and *R*^2^ on the state variables, focusing on *P/X*. In addition, we perform cross-validation using the leave-one-out strategy (predicting one process using all other processes) and compare *R*^2^ for this as well. As an alternative metric for the goodness of the model, we calculate the corrected Akaike information criterion AIC_C_[17]

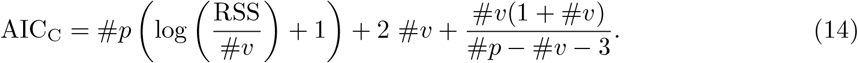

RSM only predicts end points of processes with constant exponential feed rates (*µ*_*f*_) and constant temperatures (*T*). To ensure a fair comparison, here, we restricted OptFed to the same constraints.

In addition, we validate the OptFed framework by applying it to a case study and experimentally test the predicted optimal controls.

### 2.4. Case study

We illustrate our modeling framework by optimizing protein L production in a fedbatch fermentation of *E. coli*, and evaluate its effectiveness in comparison to RSM in central composite design [10]. Data from twelve fermentations with varying specific substrate feed and temperature [51] are used as training input for both methods to predict an optimal bioprocess that maximizes the specific process yield 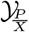

The dataset represents nine conditions (Figure 2a):

**Figure 2:**
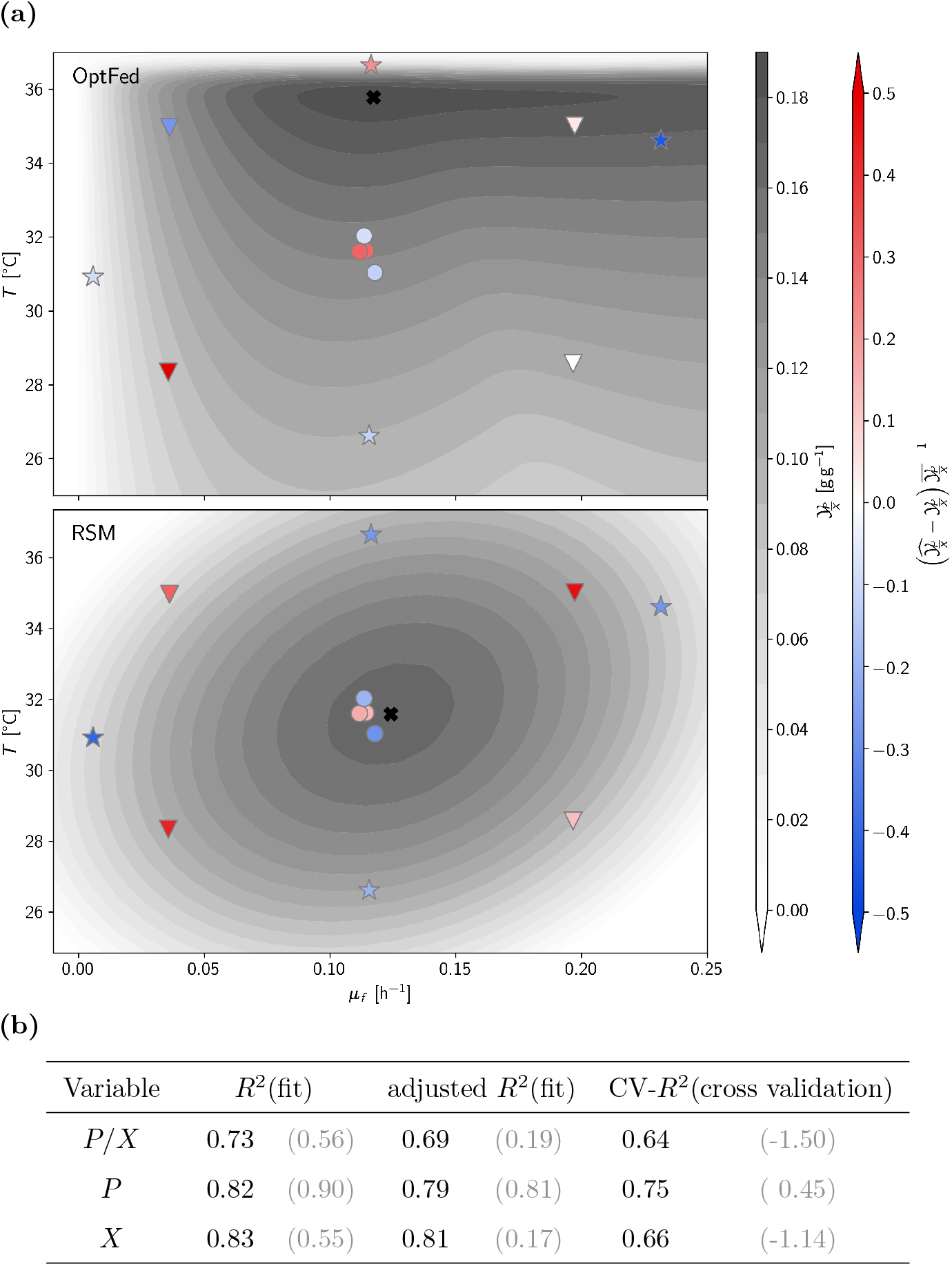
Estimation of final product titer and model errors. (a) predicted specific yield in OptFed (top) versus RSM (bottom). Both models are constructed using data from twelve fermentations, including a center point (circle, four replicates), four factorial points (triangles), and four star points (stars; each single runs). The predicted specific product yields 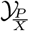 at the end of the process are shown in shades of gray, with the black cross indicating the optima. Both optima are calculated for exponential feed rates at constant temperature throughout the production phase. Model errors, indicating differences between predicted and measured 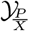, are indicated by the color of the markers: red for overestimation, white for accurate prediction, and blue for underestimation. (b) Goodness of fit [*R*^2^(fit), and adjusted *R*^2^(fit)] as well as goodness of fit for leaveone-out cross-validation (CV) [as measured by CV-*R*^2^(cross validation)]. Values in brackets refer to the RSM model, values in standard print to OptFed.

- one center point (four runs),
- four star points (single runs) where either feeding rate or temperature varied from the center point,
- four factorial points (single runs) where both variables deviated from the center point.

#### Training Data

Process data (biomass concentration, protein L concentration, and substrate concentration over time) from 12 fed-batch fermentations of *E. coli* strain BL21 DE3, with varying specific substrate feed and temperature [51], are used to fit the general process model.

In short, protein L accumulated intracellularly, and glycerol was the sole carbon source. IPTG (isopropyl *β*-D-1-thiogalactopyranoside) was used to induce the product promoter during the feed phase. The biomass concentration at induction varied between 20 g L^−1^ to 45 g L^−1^. The control variables were the exponential feed rate coefficient (*µ*_*f*_) and the temperature (*T*). Each process had a production phase of 12 h. Biomass and product concentrations were measured every 2 h, while temperature and feed rate were measured online. Due to the small reactor size, each sampling reduced the reactor volume non-negligibly, which was considered in the analysis. Details about the experimental setup and analytical methods are described in Section 2.4.1.

##### 2.4.1 Experimental procedures

Experimental data for model fitting and validation in OptFed was obtained from 15 bioreactor cultivations. The cultivations were carried out in two bioreactor systems having a similar working volume. The 12 initial cultivations were executed in a DASGIP^©^ Parallel Bioreactor System (max. working volume 2 L; Eppendorf, Hamburg, Germany) as described in [51]. In contrast to the original paper, where 11 out of 12 performed processes were used, according to the DoE, we use all 12 processes as training data. In addition, three validation runs were performed in a Minifors 2 bioreactor system (max. working volume 2.5 L; Infors HT, Bottmingen, Switzerland). For all cultivations a defined minimal medium according to DeLisa [52] was used, supplemented with an initial concentration of 20 g L^−1^ glycerol as the main carbon source and 0.02 g L^−1^ kanamycin as a selection marker. The temperature was set to 37 °C during batch phase, 35 °C during fed-batch phase and controlled at defined levels during induced fed-batch phase in accordance with the experimental plan. The pH was monitored with an EasyFerm pH electrode (Hamilton, Reno, NV, USA) and kept constant at 6.7 via addition of 12.5% NH_4_OH. A probe for monitoring dissolved oxygen (dO_2_) was installed (Visiferm DO425, Hamilton, Reno, NV, USA). The dissolved oxygen in the cell broth was kept over 40% through continuous stirring (1400 rpm) and aeration of 2 vvm. If needed, pure oxygen was added to the air flow. Furthermore, the off-gas was analyzed with respect to O_2_ and CO_2_ concentrations via a Bluevary sensor (BlueSens Gas analytics, Herten, Germany) for realtime monitoring of the metabolic activity of the cells. The process parameters were logged and controlled using the bioprocess management system eve^©^ (Infors HT, Bottmingen, Switzerland). The expression of recombinant protein L was induced by a one-point addition of sterile Isopropyl *β*-D-1-thiogalactopyranoside (IPTG) to a final concentration of 0.5 mM. After addition of the inducer, samples were taken every two hours for further process and product analytics.

An *E. coli* BL21 (DE3) strain transformed with a pET-24a(+) plasmid was used for the cultivations (GenBank accession no. AAA67503). The plasmid carries the codon-optimized genes coding the 5B (binding) protein L with a C-terminal His_6_-tag. The recombinant protein L is expressed intracellularly. The cells were harvested and subsequent analytics were done. All subsequent analytical steps were realised with samples of 35 mL cell broth each. The cell broth was centrifuged (10 min, 21 000 rpm, 4 °C) and the supernatant was separated from the cell pellet and aliquoted (1 mL) for anion exchange chromatography. Biomass concentration was quantified by dry cell weight (DCW) in triplicates. Therefore, the cell pellet was washed with saline (0.9 wt.% NaCl), centrifuged with the same settings and dried at 105 °C for 48 h. In addition, the biomass concentration was determined via optical density measurements at 600 nm wavelength (OD600) in triplicates. Residual glycerol and metabolites in the cell-free supernatant were analyzed by a high performance liquid chromatography (HPLC) system (UltiMate 3000; Thermo Fisher, Waltham, MA) equipped with an Aminex HPX-87 H column (Bio-Rad Laboratories, Hercules, CA, USA). HPLC standards with various concentrations of protein L (0.063-1.0 g L^−1^), glycerol (0.781-50 g L^−1^) and acetate (1-10 g L^−1^) were prepared separately. A sample volume of 10 mL cell broth was centrifuged (15 min, 14 000 rpm, 4°C) and the separated cell pellet was re-suspended in 40 mL lysis buffer (10 mM EDTA, 100 mM Tris, pH 7.4) and homogenized subsequently (7 passages, 1200 bar; PandaPLUS, Gea AG, Germany). After centrifugation of the crude cell lysate (20 min, 14 000 rpm, 4 °C), the supernatant was analyzed using a reversed-phase HPLC method for protein L quantification based on a PpL standard calibration curve. The UltiMate 3000 HPLC system was equipped with a BioResolve reversed-phase Polyphenyl column (Waters Corporation, MA, USA). Further information about the analytical procedures can be found in [51].

## 3. Results

We illustrate our modeling framework by optimizing protein L production in a fedbatch fermentation of *E. coli*, and evaluate its effectiveness in comparison to RSM [49] in central composite design [10]. Data from twelve fermentations with varying specific substrate feed and temperature [51] are used as input for both methods to predict an optimal bioprocess that maximizes the specific process yield 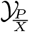 (product per biomass ratio at the end of the process).

### 3.1. Response surface methodology predicts limited optimization potential

RSM utilizes a second-degree polynomial model to forecast the specific yield 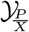 as a function of the specific feeding rate and temperature (Equation 11). It predicts an optimal specific yield of 0.16 g g^−1^ near the center point at *µ*_*f*_ = 0.12 h^*−*1^ and *T* = 31 ^*°*^C (Figure 2a). However, the predicted yield improvement is small (+1 %) yet uncertain (adjusted *R*^2^ = 0.19; Figure 2b), and none of the model’s parameters are statistically significant (at a 0.95 confidence level, Supplementary Table S3).

### 3.2. OptFed identifies significant optimization potential at high temperature

#### 3.2.1. Model Simplification and Parameter Estimation

Initially, we mitigate measurement errors of the state variables by fitting them with cubic splines (Supplementary Figure S1). Based on these splines, we parameterize our general model using the algorithm described in the Methods Section 2.1.2. We found that only 12 out of 27 parameters of the general process model are statistically significant and required (*α* = 0.2, Supplementary Figure S2), with only a minor drop in the explained variance (Table 2). The difference of *R*^2^ and adjusted *R*^2^ is higher for the full model as the degrees of freedom are higher. The effect is especially pronounced for *γ*^*°*^ as only 3 out of 12 processes are used for estimation (for other processes the substrate concentration is below the limit of quantification). Further reducing the model would increase the fitting error by at least one-third (Table 1). Thus, with the parameter values listed in Table 1, the reduced model reads:

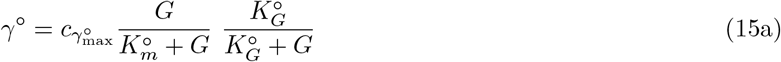

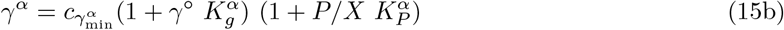

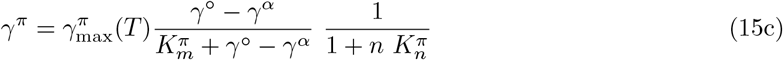

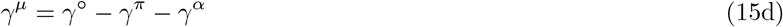

with

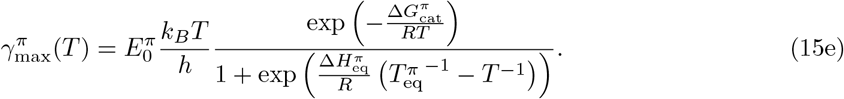

In the reduced OptFed process model (15), the substrate uptake rate *γ*^*°*^ follows Michaelis-Menten kinetics with self-inhibition by the substrate. The substrate-tomaintenance flux *γ*^*α*^ increases multilinearly with the substrate uptake rate *γ*^*°*^, and the product-to-biomass yield *P/X*. While both fluxes are temperature-independent, the product formation rate *π* is modeled as a temperature-dependent MichaelisMenten-like kinetic with non-competitive inhibition by the number of generations *n* after induction. Supplementary Figure S3 illustrates the quality of our model’s fit on rates. Despite some noise and occasional large errors in individual data points, the overall trend is well-predicted.

**Table 1:**
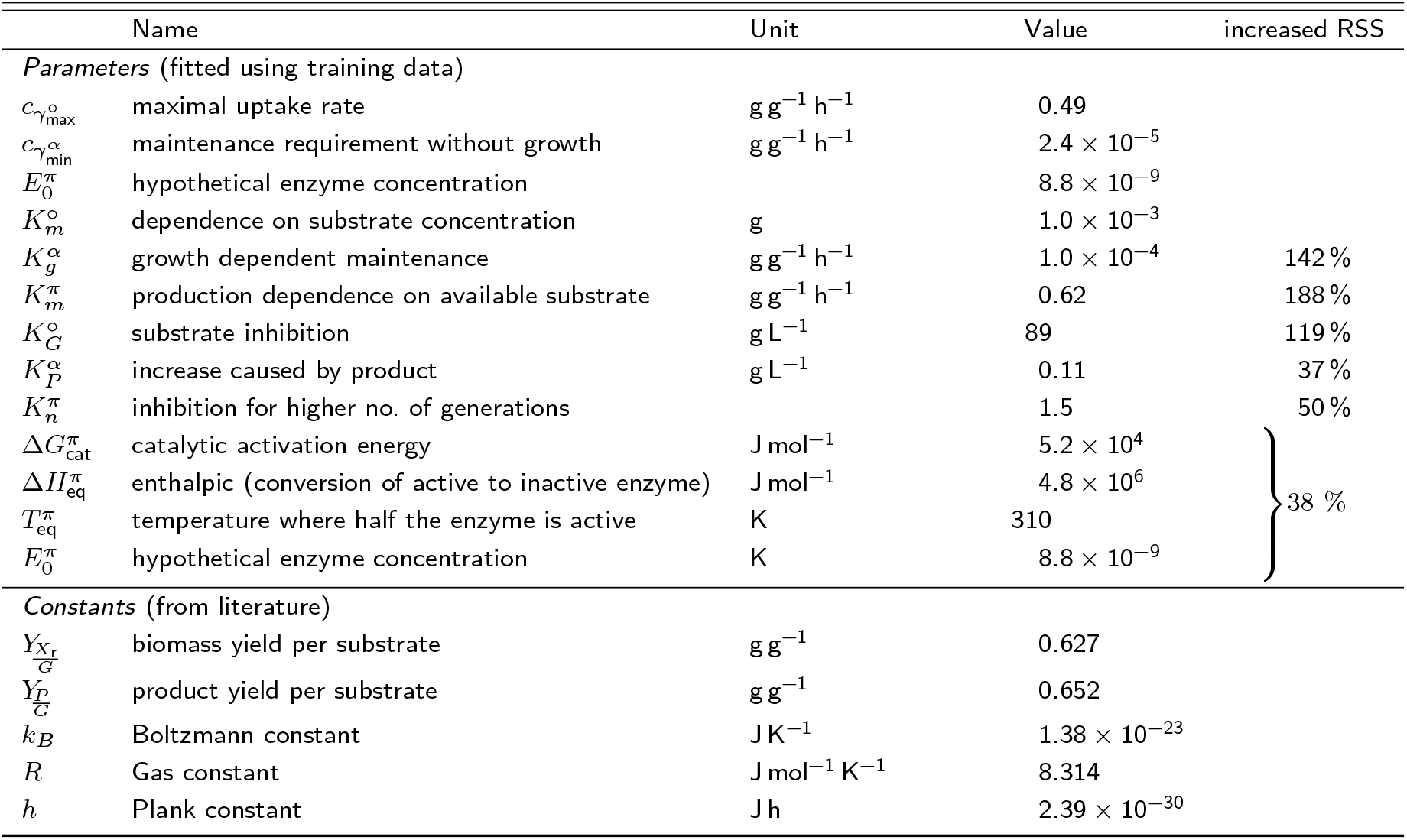
List of parameters remaining in the selected model and their fitted values. The increased RSS (residual sum of squares) column next to a parameter shows the increase in fitting error (calculated with Equation 12) if this parameter would be removed from the reduced model. *Km°* is not removed, as removal leads to physically impossible negative substrate concentrations. A comprehensive list of all parameters of the initially designed model is found in Supplementary Table S1).

**Table 2:** Overview of the goodness of fit of the kinetic functions. The values of the parameters of the reduced OptFed are given in Table 1.

|  | reduced OptFed |  |  | full OptFed |  |  |
| --- | --- | --- | --- | --- | --- | --- |
| | $\gamma^\circ$ | $\gamma^\alpha$ | $\gamma^\pi$ | $\gamma^\circ$ | $\gamma^\alpha$ | $\gamma^\pi$ |
| $R^2$ | 0.54 | 0.68 | 0.70 | 0.58 | 0.74 | 0.73 |
| adjusted $R^2$ | 0.52 | 0.67 | 0.68 | 0.24 | 0.71 | 0.70 |
| # of parameters | 3 | 3 | 6 | 9 | 9 | 9 |

We compare the measurement data for product and biomass with the model predictions and validate these predictions using cross-validation. The adjusted *R*^2^ values remain above 0.52 for *P, P/X*, and *X*, both with and without cross-validation (Figure 2b and Supplementary Figures S4 and S5). Thus, we conclude that the model is a reliable choice, especially compared to RSM.

Additionally, we compare our heuristic model selection algorithm (Section 2.1.2) with model selection based on the AIC_C_. The selected kinetic parameters are almost identical for both methods, only AIC_C_ includes terms for *G* in *γ*^*α*^ and *γ*^*π*^. With our heuristic algorithm, these variables were removed in the last step of our elimination as they do not improve the model fit significantly (*p*-values of 0.25 and 0.30). While optimal feed rates are different (about ±22 %), temperature optimum is similar (±0.03 °C). Using the model optimum for one model and testing it with the other misses the optimal 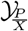 by less than 0.6 %.

#### 3.2.2. Process Model Optimization

Using the model develop above, we apply the optimization algorithm described in Methods Section 2.1.3 to determine the optimal feeding strategy and temperature settings. Additionally, we consider the following constraints:

- The initial biomass matches the mean value of the center point runs of the experimental data, *X*_0_ = 30 g L^*−*1^.
- The initial bioreactor volume is set at 1.3 L, with no maximum volume constraint.
- The feed glycerol concentration is fixed at 390 g L^−1^.
- Sampling of 35 mL reduces the current reactor volume at *t* = 2, 4, 6, 8, and 10 h.
- A linear regression model (Supplementary Figure S6) approximates the base addition as

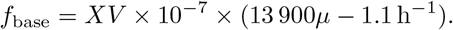

Figure 3 illustrates the predicted optimal fed-batch process for protein L production. At 35.8 °C and after 12 h, we predict a biomass concentration of 52 g L^−1^ and a product concentration of 9.6 g L^−1^, resulting in an optimal product yield of 0.19 g g^−1^. This represents a 37.1 % increase compared to the reference process.

**Figure 3:**
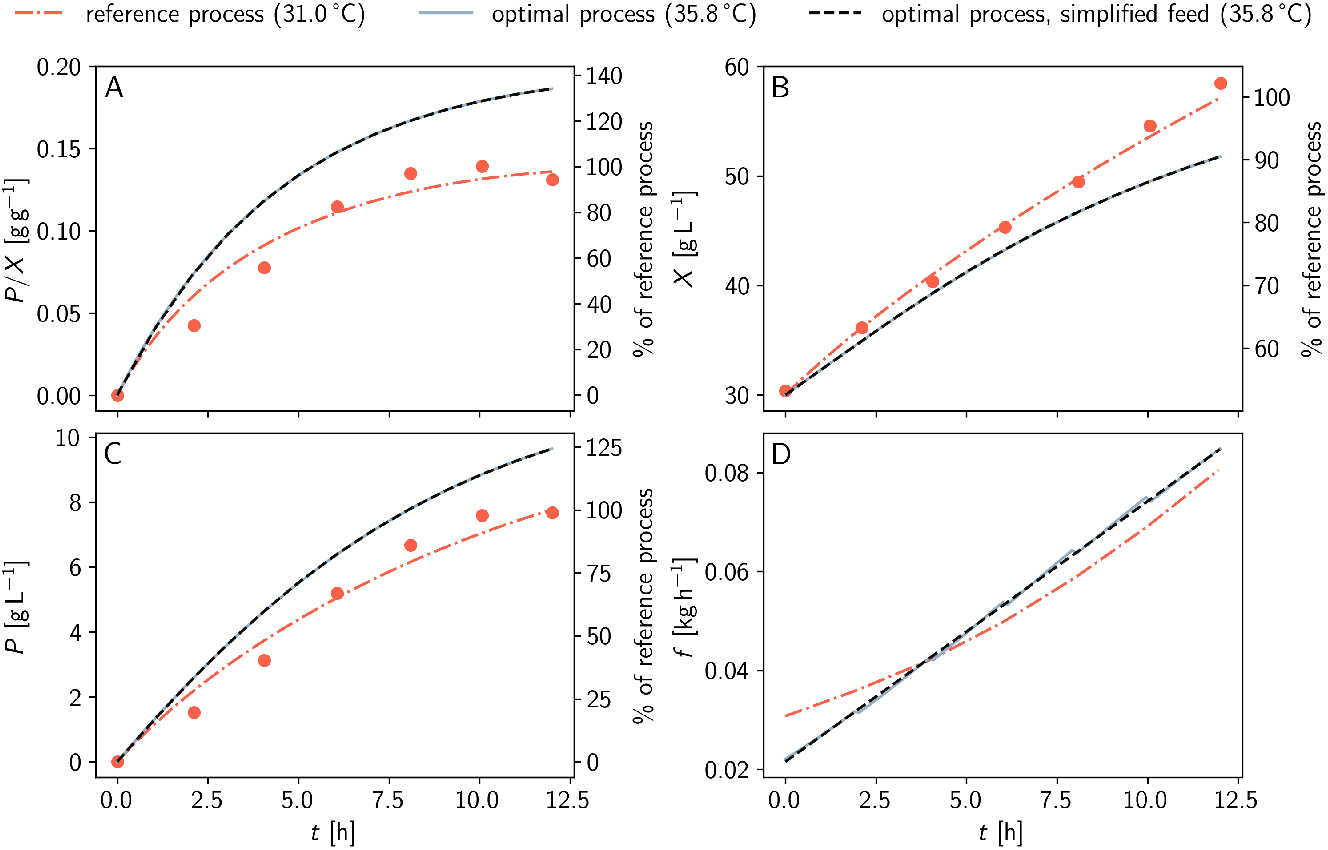
Predicted optimal process with OptFed. The red dashed-dotted lines represent OptFed’s prediction for the reference process, while grey lines show the predicted optimal behavior at *T* = 35.8 *°*C. Black dashed lines indicate the optimal process with approximated linear feed rate. The simplification of the feed profile has a negligible impact on the process (grey full and black dashed lines). After 12 h, a 37.4 % improvement in specific product yield is predicted. For reference, measurements from the training data are shown (red circles, center point, same initial biomass).

Increasing the temperature is key for optimized performance (Figure 3). According to the model, maximum production rate 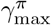 increases with temperature up to the optimal temperature of 35.8 °C and decreases sharply for higher temperatures (Supplementary Figure S3). Optimizing the feed but keeping the temperature at 31 °C increases the specific product yield by just 0.3 %. Conversely, keeping the feed constant and raising only the temperature boosts the maximally obtainable specific product yield by 37.0 %. A summary of optimization results can be found in Supplementary Table S4.

Figure 2a compares the predictions of RSM and OptFed. Under identical process constraints (constant exponential feed rate and constant temperature), OptFed identifies an optimum at elevated temperatures that RSM misses.

Unlike the training data’s fermentations, which used an exponential feed, we find that an almost linearly increasing feed rate is best to maximize the product-to-biomass yield (Figure 3D). Therefore, we decided to approximate the predicted optimal feed function with a simple linear equation (Supplementary Figure S6)

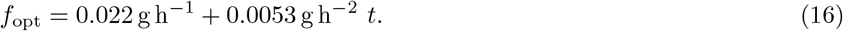

This adjustment changes the final product and biomass concentrations and the product-to-biomass yield 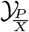 by less than 0.1 %, but significantly eases practical implementation (Figure 4). Generally, variations of the feed function have little influence as long as the initial and final biomass concentrations remain constant.

**Figure 4:**
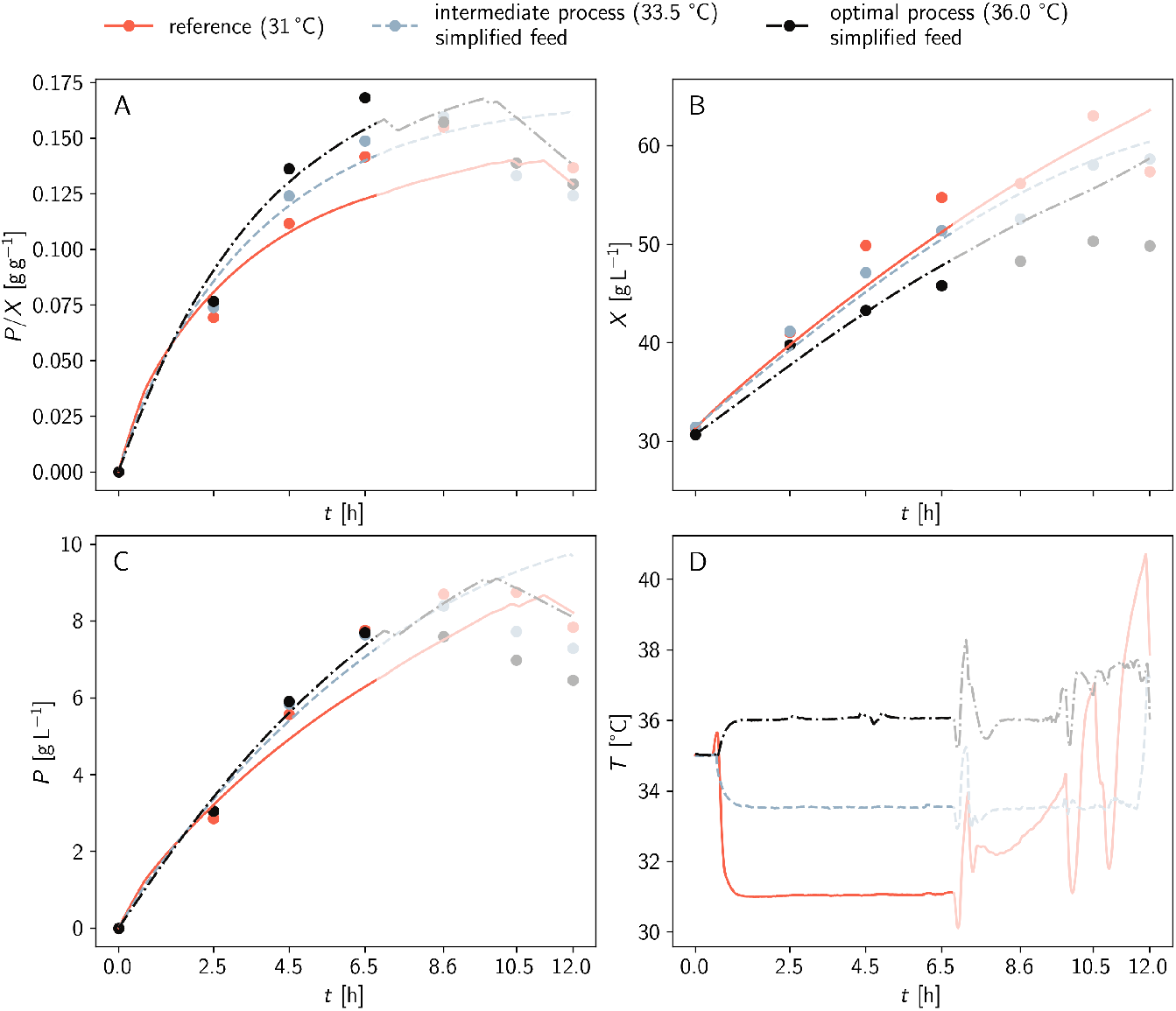
Validation of the predicted optimal fermentation. Panels A to C compare experimental data (circles) with model predictions (lines). Corresponding feed rates, substrate uptake rates, growth rates, and product formation rates are illustrated in Supplementary Figure S9. Panel D shows the temperature profile of the fermentations. For *t >* 6.5 h temperatures are unstable and the data is not considered. (opaque region in panels A to D). A comparison of optimal predicted and experimentally achieved controls is given in Supplementary Figure S7.

Our model’s predictions are validated by experimentally running the optimal fermentation process with the simplified linear feed (Figure 4). Additionally, an intermediate process at 33.5 °C (halfway between the temperature of the center point and the optimum) is carried out.

For the first 6.5 h, the processes closely matched the predictions (Figure 4). The optimal process at 36 °C achieved a 19 % increase in specific product yield (compared to the predicted 21 %), while the process at 33.5 °C achieved a 5 % increase (compared to the predicted 18 %). Data points beyond 6.5 h are affected by unstable temperatures and therefore not considered.

## 4. Discussion

Recent years have seen a surge in research on biotechnological process parameterization and optimization strategies [21, 53–55]. However, studies usually focus on one of the two aspects. Here, we developed a comprehensive modeling framework, OptFed, to strategically combine them.

OptFed employs a general phenomenological process model fitted with experimental time series data from multiple fermentations. The algorithm discards parameters with insufficient statistical power to minimize overfitting, simplify the model, and increase the model’s reliability. Using this parameterized model, we applied nonlinear optimization to predict a fermentation profile for optimal specific protein L yield in *E. coli*, resulting in a near-linear feed at an elevated temperature of 35.8 °C. This approach predicted a 37 % increase in specific product yield compared to the training data, significantly outperforming the standard RSM, which only predicted a 1 % improvement. However, during the experimental validation of the optimized process, we encountered issues with temperature stability shortly after 6.5 h and decided not to use data collected thereafter. This instability is due to an undersized cooling capacity of the bioreactor.

Despite this, at 6.5 h the specific protein yield was up by 19 %, close to the predicted 21 % at that time. The primary increase in specific product yield is attributed to the elevated temperature, accounting for over 99 % of the improvement. According to OptFed, product concentration increases approximately linearly with temperature up to the optimum. This dependence is completely missed by RSM, highlighting the advantage of our approach.

Compared to RSM, our process equations constrain the possible solution space to more realistic outcomes. In fact, combining mechanistic modeling with purely statistical approaches has already previously been shown to perform better than pure RSM [54]. For example, RSM may predict negative values of the target metric (Figure 2), an effect that cannot be observed with OptFed. Moreover, our process equations implicitly ensure mass balance due to the calculation of the substrate-to-growth rate (*γ*^*µ*^) from the difference of substrate uptake and the other substrate draining fluxes (Equation 2).

Maintenance flux, as we use it throughout the manuscript, is defined as a catchall-term for several metabolic effects. It comprises (1) the (non-)growth associated maintenance [56], (2) all non-optimal growth and production due to byproduct formation [57], and (3) any overflow metabolism during high substrate uptake rates [58]. Consequently, our yields are derived from a metabolomic model, excluding maintenance requirements, which differs from experimental yields where maintenance is included. Maintenance accounts for more than half of the uptake at the end of production (Supplementary Figure S8), leading to seemingly higher-thanusual yields 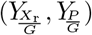 in Table 1.

Furthermore, we estimate uptake rates during production based on experimental data. As uptake can be significantly reduced during production [59] this avoids possible overfeeding in the predicted optimum. In the case study, we observe a reduced uptake rate, but uptake is not a limiting factor for optimization.

The mathematical problems in our procedure, such as model identification and calibration, are difficult to solve because models often have too many interrelated parameters and lack sufficient high-quality training data. This can cause optimization algorithms to be unstable and not converge, presenting significant challenges [60]. Our model, for example, requires high-quality time course data. Random (relative) errors for biomass and product concentrations should be in the range of 3 % and 15 %, respectively, to reliably identify the correct optimum (Supplementary Figure S10). However, these experimental uncertainties are manageable with current process monitoring technology [61].

Compared to the center point of the training data, almost all of the improvement in our case study originates from the increase in temperature. However, due to different product formation kinetics, this may be different for other products. Therefore, we cannot derive a general rule which feed profiles and temperatures are more advantageous in other setups. For example, [62] find a high influence of the feeding strategy on the production of inclusion bodies. This is most likely to a difference in product and process setup.

In Section 2.1.3, we optimize for the product-to-biomass yield 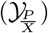, a rather unconventional metric, compared to the titer, productivity, and (product-to-substrate) yield commonly used [63]. However, in this case study, the maximization of 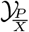 is of critical importance for the ease of downstream processing. This strategy bears the risk of converging to a process that yields excellent 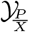 but very low amounts of product overall, which is also unfavorable. Here, we mitigated the risk by comparing our optimal 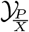 process to an optimal *PV* process (Supplementary Figure S11). Volume can be easily scaled by scaling batch volume and feed rate. The reachable biomass concentration depends on the reactor design (cooling capabilities, oxygen supply). A biomass limit of 60 g L^−1^ (reached in training processes) could increase the product by 16 %.

While our optimum is stable for changes in feeding strategy, we see a sharp drop in productivity when the temperature is raised above the optimum. This means, the optimal temperature is a good first guess, but more data is required when we move from screening to the design of the production setup. Based on the existing data, we applied a Monte Carlo estimation [64–66] (Supplementary Figure S12). We observe that we can guarantee (at a significance level of 0.05) a production rate within 10 % of the optimum by reducing the temperature from 35.8 °C to 34.9 °C.

OptFed focuses on the model selection. Based on this further improvements are possible. Sensitivity analyses [67] could help to make the optimum more stable considering the uncertainties in parameter fitting. We also limit ourselves by using existing data. Model-based design of experiment [68] could provide better training data or can be used to plan additional experiments to improve the model.

## 5. Conclusion

In this study, we presented OptFed, a phenomenological model-based bioprocess optimization framework that (also) allows us to seamlessly integrate preexisting biological and process knowledge. Unlike other tools, we emphasized the parameterization of process equations using experimental bioprocess data. To prevent overfitting, OptFed employs a multi-step fitting strategy that retains only the terms that significantly reduce model error, discarding others.

This approach addresses key challenges in industrial process design by eliminating the reliance on trial-and-error methods and standard, predefined feeding strategies. We demonstrated that OptFed not only accurately describes the training data but also predicts optimized process controls. Experimental validation shows a 19 % increase in specific protein L yield compared to the control.

While effective, OptFed’s performance depends on the quality of the training data. Future work will explore expanding the model to address more complex biological phenomena, incorporate multi-objective optimization, and validate its application across a broader range of bioprocesses and products.

We are confident that OptFed is a valuable tool for bioprocess optimization and will benefit the industry in the future.

## Competing interests

The authors declare that they have no competing interests.

## Author’s contributions

GS: conceptualization, methodology, software, investigation, visualization, writing – original draft, review and editing; RL: investigation, writing – review and editing; SK: investigation, writing – review and editing; OS: funding, writing – review and editing;, JK: investigation, writing – review and editing; JZ: funding, methodology, writing – original draft, review and editing; MG: methodology, writing – original draft, review and editing.

## Author details

^1^ Department of Analytical Chemistry, University Vienna, Währinger Straße, 1090 Vienna, Austria, EU. ^2^ Doctorate School of Chemistry, University of Vienna, Währinger Straße, 1090 Vienna, Austria, EU. ^3^ Integrated Bioprocess Development, Technical University Vienna, Getreidemarkt 9, 1060 Vienna, Austria, EU. ^4^ Austrian Centre of Industrial Biotechnology, Krenngasse 37, 8010 Graz, Austria, EU.

## Notes

https://github.com/gschloegel/OptFed

### Competing Interest Statement

The authors have declared no competing interest.

### Summary of Updates

Updated all sections according to reviewers' comments.

## References

1. Henry C Lim and Hwa Sung Shin. Fed-batch cultures: principles and applications of semi-batch bioreactors. Cambridge University Press, 2013.

2. Maria Isabel Rodrigues and Antonio Francisco Iemma. Experimental Design and Process Optimization. CRC Press, Boca Raton, December 2014. ISBN 978-0-429-16186-5..

3. JM Modak, HC Lim, and YJ Tayeb. General characteristics of optimal feed rate profiles for various fed-batch fermentation processes. Biotechnology and bioengineering, 28(9):1396–1407, 1986.

4. Michael Maurer, Manfred Kühleitner, Brigitte Gasser, and Diethard Mattanovich. Versatile modeling and optimization of fed batch processes for the production of secreted heterologous proteins with pichia pastoris. Microbial cell factories, 5(1):1–10, 2006.

5. David M. Steinberg and Dizza Bursztyn. Response Surface Methodology in Biotechnology. Quality Engineering, 22(2):78–87, March 2010. ISSN 0898-2112. URL 10.1080/08982110903510388. Publisher: Taylor & Francis eprint: 10.1080/08982110903510388.

6. Rafael D de Oliveira, Galo AC Le Roux, and Radhakrishnan Mahadevan. Nonlinear programming reformulation of dynamic flux balance analysis models. Computers & Chemical Engineering, 170:108101, 2023.

7. Steffen Klamt, Radhakrishnan Mahadevan, and Oliver Hädicke. When do two-stage processes outperform one-stage processes? Biotechnology journal, 13(2):1700539, 2018.

8. Grégory Mermoud. Model-Based Optimization. In Gregory Mermoud, editor, Stochastic Reactive Distributed Robotic Systems: Design, Modeling and Optimization, Springer Tracts in Advanced Robotics, pages 175–179. Springer International Publishing, Cham, 2014. ISBN 978-3-319-02609-1. URL 10.1007/978-3-319-02609-1_11.

9. João C. M. Carvalho, Michele Vitolo, Sunao Sato, and Eugenio Aquarone. Ethanol production by Saccharomyces cerevisiae grown in sugarcane blackstrap molasses through a fed-batch process. Applied Biochemistry and Biotechnology, 110(3):151–164, September 2003. ISSN 1559-0291. URL 10.1385/ABAB:110:3:151.

10. André I. Khuri and Siuli Mukhopadhyay. Response surface methodology. WIREs Computational Statistics, 2(2):128–149, 2010. ISSN 1939-0068. URL http://onlinelibrary.wiley.com/doi/abs/10.1002/wics.73. eprint: https://wires.onlinelibrary.wiley.com/doi/pdf/10.1002/wics.73.

11. Jacques Monod. The growth of bacterial cultures. Annual Review of Microbiology, 3(1):371–394, October 1949. ISSN 0066-4227. URL https://www.annualreviews.org/doi/10.1146/annurev.mi.03.100149.002103. Publisher: Annual Reviews.

12. Marta B. Lopes, Gabriel Martins, and Cecília R. C. Calado. Kinetic modeling of plasmid bioproduction in Escherichia coli DH5α cultures over different carbon-source compositions. Journal of Biotechnology, 186: 38–48, September 2014. ISSN 0168-1656. URL https://www.sciencedirect.com/science/article/pii/S0168165614003137.

13. Stefan Klumpp, Zhongge Zhang, and Terence Hwa. Growth Rate-Dependent Global Effects on Gene Expression in Bacteria. Cell, 139(7):1366–1375, December 2009. ISSN 0092-8674, 1097-4172. URL https://www.cell.com/cell/abstract/S0092-8674(09)01505-0. Publisher: Elsevier.

14. Julian Kager, Johanna Bartlechner, Christoph Herwig, and Stefan Jakubek. Direct control of recombinant protein production rates in E. coli fed-batch processes by nonlinear feedback linearization. Chemical Engineering Research and Design, 182:290–304, June 2022. ISSN 0263-8762. URL https://www.sciencedirect.com/science/article/pii/S0263876222001460.

15. Jan Weber, Frank Hoffmann, and Ursula Rinas. Metabolic adaptation of Escherichia coli during temperature-induced recombinant protein production: 2. Redirection of metabolic fluxes. Biotechnology and Bioengineering, 80(3):320–330, 2002. ISSN 1097-0290. URL https://onlinelibrary.wiley.com/doi/abs/10.1002/bit.10380. eprint: https://onlinelibrary.wiley.com/doi/pdf/10.1002/bit.10380.

16. Holger W. Jannasch and Thomas Egli. Microbial growth kinetics: a historical perspective. Antonie van Leeuwenhoek, 63(3):213–224, September 1993. ISSN 1572-9699. URL 10.1007/BF00871219.

17. CLIFFORD M. Hurvich and CHIH-LING Tsai. Regression and time series model selection in small samples. Biometrika, 76(2):297–307, June 1989. ISSN 0006-3444. URL 10.1093/biomet/76.2.297.

18. Jason D. Lee, Dennis L. Sun, Yuekai Sun, and Jonathan E. Taylor. Exact post-selection inference, with application to the lasso. The Annals of Statistics, 44(3):907–927, June 2016. ISSN 0090-5364, 2168-8966. URL https://projecteuclid.org/journals/annals-of-statistics/volume-44/issue-3/Exact-post-selection-inference-with-application-to-the-lasso/10.1214/15-AOS1371.full. Publisher: Institute of Mathematical Statistics.

19. Yongli Zhang and Yuhong Yang. Cross-validation for selecting a model selection procedure. Journal of Econometrics, 187(1):95–112, July 2015. ISSN 0304-4076. URL https://www.sciencedirect.com/science/article/pii/S0304407615000305.

20. Luke A. Yates, Zach Aandahl, Shane A. Richards, and Barry W. Brook. Cross validation for model selection: A review with examples from ecology. Ecological Monographs, 93(1):e1557, 2023. ISSN 1557-7015. URL https://onlinelibrary.wiley.com/doi/abs/10.1002/ecm.1557. eprint: https://onlinelibrary.wiley.com/doi/pdf/10.1002/ecm.1557.

21. Benjamín J. Sánchez, Daniela C. Soto, Héctor Jorquera, Claudio A. Gelmi, and José R. Pérez-Correa. HIPPO: An Iterative Reparametrization Method for Identification and Calibration of Dynamic Bioreactor Models of Complex Processes. Industrial & Engineering Chemistry Research, 53(48):18514–18525, December 2014. ISSN 0888-5885. URL 10.1021/ie501298b. Publisher: American Chemical Society.

22. Khuloud Jaqaman and Gaudenz Danuser. Linking data to models: data regression. Nature Reviews Molecular Cell Biology, 7(11):813–819, November 2006. ISSN 1471-0080. URL https://www.nature.com/articles/nrm2030. Publisher: Nature Publishing Group.

23. Sudhir Varma and Richard Simon. Bias in error estimation when using cross-validation for model selection. BMC Bioinformatics, 7(1):91, February 2006. ISSN 1471-2105. URL 10.1186/1471-2105-7-91.

24. Gerd Gigerenzer and Henry Brighton. Homo Heuristicus: Why Biased Minds Make Better Inferences. Topics in Cognitive Science, 1(1):107–143, 2009. ISSN 1756-8765. URL https://onlinelibrary.wiley.com/doi/abs/10.1111/j.1756-8765.2008.01006.x. eprint: https://onlinelibrary.wiley.com/doi/pdf/10.1111/j.1756-8765.2008.01006.x.

25. Douglas M. Hawkins. The Problem of Overfitting. Journal of Chemical Information and Computer Sciences, 44 (1):1–12, January 2004. ISSN 0095-2338. URL 10.1021/ci0342472. Publisher: American Chemical Society.

26. Jens Nielsen and Jay D Keasling. Engineering cellular metabolism. Cell, 164(6):1185–1197, 2016.

27. Jorge Nocedal and Stephen J. Wright. Numerical optimization. Springer series in operations research. Springer, New York, 2nd ed edition, 2006. ISBN 978-0-387-30303-1. OCLC: ocm68629100.

28. Dante Kalise, Karl Kunisch, and Zhiping Rao. Hamilton-Jacobi-Bellman Equations: Numerical Methods and Applications in Optimal Control. De Gruyter, August 2018. ISBN 978-3-11-054359-9. URL http://www.degruyter.com/document/doi/10.1515/9783110543599/html. Publication Title: Hamilton-Jacobi-Bellman Equations.

29. Daniel Liberzon. Calculus of Variations and Optimal Control Theory. Princeton University Press, 2012. ISBN 978-0-691-15187-8.

30. B. Srinivasan, S. Palanki, and D. Bonvin. Dynamic optimization of batch processes: I. Characterization of the nominal solution. Computers & Chemical Engineering, 27(1):1–26, January 2003. ISSN 0098-1354. URL https://www.sciencedirect.com/science/article/pii/S0098135402001163.

31. V. S. Vassiliadis, R. W. H. Sargent, and C. C. Pantelides. Solution of a Class of Multistage Dynamic Optimization Problems. 1. Problems without Path Constraints. Industrial & Engineering Chemistry Research, 33(9):2111–2122, September 1994. ISSN 0888-5885. URL 10.1021/ie00033a014. Publisher: American Chemical Society.

32. Richard E. Bellman. Dynamic Programming. Princeton University Press, August 2021. ISBN 978-1-4008-3538-6. URL http://www.degruyter.com/document/doi/10.1515/9781400835386/html. Publication Title: Dynamic Programming.

33. Bojan Bojkov and Rein Luus. Time-Optimal Control by Iterative Dynamic Programming. Industrial & Engineering Chemistry Research, 33(6):1486–1492, June 1994. ISSN 0888-5885. URL 10.1021/ie00030a008. Publisher: American Chemical Society.

34. Joel A. E. Andersson, Joris Gillis, Greg Horn, James B. Rawlings, and Moritz Diehl. CasADi: a software framework for nonlinear optimization and optimal control. Mathematical Programming Computation, 11(1): 1–36, March 2019. ISSN 1867-2957. URL 10.1007/s12532-018-0139-4.

35. Logan D. R. Beal, Daniel C. Hill, R. Abraham Martin, and John D. Hedengren. GEKKO Optimization Suite. Processes, 6(8):106, August 2018. ISSN 2227-9717. URL https://www.mdpi.com/2227-9717/6/8/106. Number: 8 Publisher: Multidisciplinary Digital Publishing Institute.

36. William E. Hart, Jean-Paul Watson, and David L. Woodruff. Pyomo: modeling and solving mathematical programs in Python. Mathematical Programming Computation, 3(3):219, August 2011. ISSN 1867-2957. URL 10.1007/s12532-011-0026-8.

37. Jingyi Yang, Yuebao Yang, and Mingtao Li. OptControl.jl: An Interpreter for Optimal Control Problem, July 2022. URL http://arxiv.org/abs/2207.13229. arXiv:2207.13229 [math].

38. Leonor Michaelis and Maud Leonora Menten. Die Kinetik der Invertinwirkung. Biochemische Zeitschrift, 49: 333–369, 1913. ISSN 0366-0753. URL http://publikationen.ub.uni-frankfurt.de/frontdoor/index/index/docId/17273.

39. Peter van Bodegom. Microbial Maintenance: A Critical Review on Its Quantification. Microbial Ecology, 53(4):513–523, May 2007. ISSN 1432-184X. URL 10.1007/s00248-006-9049-5.

40. Roy M. Daniel, Michael J. Danson, Robert Eisenthal, Charles K. Lee, and Michelle E. Peterson. The effect of temperature on enzyme activity: new insights and their implications. Extremophiles, 12(1):51–59, January 2008. ISSN 1433-4909. URL 10.1007/s00792-007-0089-7.

41. Jonathan M Monk, Anna Koza, Miguel A Campodonico, Daniel Machado, Jose Miguel Seoane, Bernhard O Palsson, Markus J Herrgård, and Adam M Feist. Multi-omics quantification of species variation of escherichia coli links molecular features with strain phenotypes. Cell systems, 3(3):238–251, 2016.

42. Gilbert F. Froment, Kenneth B. Bischoff, and Juray De Wilde. Chemical reactor analysis and design, volume 2. Wiley New York, 1990.

43. André Bardow and Wolfgang Marquardt. Incremental and simultaneous identification of reaction kinetics: methods and comparison. Chemical Engineering Science, 59(13):2673–2684, July 2004. ISSN 0009-2509. URL https://www.sciencedirect.com/science/article/pii/S0009250904002015.

44. Pauli Virtanen, Ralf Gommers, Travis E. Oliphant, Matt Haberland, Tyler Reddy, David Cournapeau, Evgeni Burovski, Pearu Peterson, Warren Weckesser, Jonathan Bright, Stéfan J. van der Walt, Matthew Brett, Joshua Wilson, K. Jarrod Millman, Nikolay Mayorov, Andrew R. J. Nelson, Eric Jones, Robert Kern, Eric Larson,J J Carey, Ilhan Polat, Yu Feng, Eric W. Moore, Jake VanderPlas, Denis Laxalde, Josef Perktold, Robert Cimrman, Ian Henriksen, E. A. Quintero, Charles R. Harris, Anne M. Archibald, Antonio H. Ribeiro, Fabian Pedregosa, Paul van Mulbregt, and SciPy 1.0 Contributors. SciPy 1.0: Fundamental Algorithms for Scientific Computing in Python. Nature Methods, 17:261–272, 2020.

45. Andrew Rutherford. Introducing Anova and Ancova : A GLM Approach. Introducing Statistical Methods. SAGE Publications Ltd, London, 2001. ISBN 978-0-7619-5160-5. URL https://search.ebscohost.com/login.aspx?direct=true&db=nlebk&AN=251737&site=ehost-live.

46. Rainer Storn and Kenneth Price. Differential Evolution – A Simple and Efficient Heuristic for global Optimization over Continuous Spaces. Journal of Global Optimization, 11(4):341–359, December 1997. ISSN 1573-2916. URL 10.1023/A:1008202821328.

47. Andreas Wächter and Lorenz T. Biegler. On the implementation of an interior-point filter line-search algorithm for large-scale nonlinear programming. Mathematical Programming, 106(1):25–57, March 2006. ISSN 1436-4646. URL 10.1007/s10107-004-0559-y.

48. Lorenz T. Biegler. Nonlinear Programming: Concepts, Algorithms, and Applications to Chemical Processes. Society for Industrial and Applied Mathematics, January 2010. ISBN 978-0-89871-702-0978-0-89871-938-3. URL http://epubs.siam.org/doi/book/10.1137/1.9780898719383.

49. Russell V. Lenth. Response-Surface Methods in R, Using rsm. Journal of Statistical Software, 32:1–17, 2010. ISSN 1548-7660. URL 10.18637/jss.v032.i07.

50. R Core Team. R: A Language and Environment for Statistical Computing. R Foundation for Statistical Computing, Vienna, Austria, 2023. URL https://www.R-project.org/.

51. Stefan Kittler, Julian Ebner, Mihail Besleaga, Johan Larsbrink, Barbara Darnhofer, Ruth Birner-Gruenberger, Silvia Schobesberger, Christopher K. Akhgar, Andreas Schwaighofer, Bernhard Lendl, and Oliver Spadiut. Recombinant Protein L: Production, Purification and Characterization of a Universal Binding Ligand. Journal of Biotechnology, 359:108–115, November 2022. ISSN 0168-1656. URL https://www.sciencedirect.com/science/article/pii/S0168165622002371.

52. Matthew P. DeLisa, Jincai Li, Govind Rao, William A. Weigand, and William E. Bentley. Monitoring GFP-operon fusion protein expression during high cell density cultivation of Escherichia coli using an on-line optical sensor. Biotechnology and Bioengineering, 65(1):54–64, 1999.

53. Jasmin Bauer and Steffen Klamt. Optmsp: A toolbox for designing optimal multi-stage (bio) processes. Journal of Biotechnology, 383:94–102, 2024.

54. José Pinto Cristiana Rodrigues de Azevedo, Rui Oliveira, and Moritz von Stosch. A bootstrap-aggregated hybrid semi-parametric modeling framework for bioprocess development. Bioprocess and biosystems engineering, 42:1853–1865, 2019.

55. Kaushik Raj, Naveen Venayak, and Radhakrishnan Mahadevan. Novel two-stage processes for optimal chemical production in microbes. Metabolic Engineering, 62:186–197, November 2020. ISSN 1096-7176. URL https://www.sciencedirect.com/science/article/pii/S1096717620301269.

56. Benjamin Luke Coltman, Corinna Rebnegger, Brigitte Gasser, and Jürgen Zanghellini. Characterising the metabolic rewiring of extremely slow growing komagataella phaffii. Microbial Biotechnology, 17(1):e14386, 2024.

57. Aristos A Aristidou, Ka-Yiu San, and George N Bennett. Improvement of biomass yield and recombinant gene expression in escherichia coli by using fructose as the primary carbon source. Biotechnology progress, 15(1):140–145, 1999.

58. Bo Xu, Mehmedalija Jahic, and Sven-Olof Enfors. Modeling of overflow metabolism in batch and fed-batch cultures of escherichiacoli. Biotechnology progress, 15(1):81–90, 1999.

59. Don Fabian Müller, Daniel Wibbing, Christoph Herwig, and Julian Kager. Simultaneous real-time estimation of maximum substrate uptake capacity and yield coefficient in induced microbial cultures. Computers & Chemical Engineering, 173:108203, May 2023. ISSN 0098-1354. URL https://www.sciencedirect.com/science/article/pii/S0098135423000728. Publisher: Pergamon.

60. Vinzenz Abt, Tilman Barz, Mariano Nicolas Cruz-Bournazou, Christoph Herwig, Paul Kroll, Johannes Möller, Ralf Pörtner, and René Schenkendorf. Model-based tools for optimal experiments in bioprocess engineering. Current opinion in chemical engineering, 22:244–252, 2018.

61. Julian Kager and Christoph Herwig. Monte Carlo-Based Error Propagation for a More Reliable Regression Analysis across Specific Rates in Bioprocesses. Bioengineering, 8(11):160, November 2021. ISSN 2306-5354. URL https://www.mdpi.com/2306-5354/8/11/160. Number: 11 Publisher: Multidisciplinary Digital Publishing Institute.

62. Christoph Slouka, Julian Kopp, Daniel Strohmer, Julian Kager, Oliver Spadiut, and Christoph Herwig. Monitoring and control strategies for inclusion body production in E. coli based on glycerol consumption. Journal of Biotechnology, 296:75–82, April 2019. ISSN 0168-1656. URL https://www.sciencedirect.com/science/article/pii/S0168165619300951.

63. Kai Zhuang, Laurence Yang, William R Cluett, and Radhakrishnan Mahadevan. Dynamic strain scanning optimization: an efficient strain design strategy for balanced yield, titer, and productivity. dyssco strategy for strain design. BMC biotechnology, 13:1–15, 2013.

64. S. T. Buckland. Monte Carlo Confidence Intervals. Biometrics, 40(3):811–817, 1984. ISSN 0006-341X. URL https://www.jstor.org/stable/2530926. Publisher: International Biometric Society.

65. Niels Krausch, Tilman Barz, Annina Sawatzki, Mathis Gruber, Sarah Kamel, Peter Neubauer, and Mariano Nicolas Cruz Bournazou. Monte Carlo Simulations for the Analysis of Non-linear Parameter Confidence Intervals in Optimal Experimental Design. Frontiers in Bioengineering and Biotechnology, 7, May 2019. ISSN 2296-4185. URL https://www.frontiersin.org/journals/bioengineering-and-biotechnology/articles/10.3389/fbioe.2019.00122/full. Publisher: Frontiers.

66. Kristopher J. Preacher and James P. Selig. Advantages of Monte Carlo Confidence Intervals for Indirect Effects. Communication Methods and Measures, 6(2):77–98, April 2012. ISSN 1931-2458. URL 10.1080/19312458.2012.679848. Publisher: Routledge eprint: 10.1080/19312458.2012.679848.

67. René Schenkendorf Xiangzhong Xie, Moritz Rehbein, Stephan Scholl, and Ulrike Krewer. The Impact of Global Sensitivities and Design Measures in Model-Based Optimal Experimental Design. Processes, 6(4):27, April 2018. ISSN 2227-9717. URL https://www.mdpi.com/2227-9717/6/4/27. Number: 4 Publisher: Multidisciplinary Digital Publishing Institute.

68. Gaia Franceschini and Sandro Macchietto. Model-based design of experiments for parameter precision: State of the art. Chemical Engineering Science, 63(19):4846–4872, October 2008. ISSN 0009-2509. URL https://www.sciencedirect.com/science/article/pii/S0009250907008871.

