## Supplementary Material for "Optimizing Bioprocessing Efficiency with OptFed: Dynamic Nonlinear Modeling Improves Product-to-Biomass Yield"

RESEARCH

### S1 Supplementery Methods

#### S1.1 Spline fit

In Section 2.1.2, the fitting of the rates $g$, $\gamma^{\mu}$, and $\gamma^{\pi}$ is described. Here we go into more detail about how they are calculated for our specific experimental setup.

The following variables are measured during the processes of the training data (see Section 2.4.1):

| | |
|---|---|
| $T$ | Temperature |
| $\widehat{V}$ | Volume (as reactor mass) |
| $f^{G}_{\text{cum}}$ | cumulative substrate feed (mass of the feed flask) |
| $f^{B}_{\text{cum}}$ | cumulative base feed for control pH (mass of the feed flask) |
| $G^{G}_{f}$ | feed concentration in the substrate feed |
| $\widehat{X}$ | biomass concentration |
| $\widehat{P}$ | product concentration |
| $\widehat{G}$ | free substrate in the media |

The first 4 values are measured continuously, and the last 3 every 2 hours.

To get continuous functions for the biomass, product, and substrate concentrations and derivatives for the cumulative feeds, spline fitting is used:

$$f = f^{\dot{G}}_{\text{cum}} + f^{\dot{B}}_{\text{cum}} \tag{S1a}$$

$$G_{\text{f}} = f^{\dot{G}}_{\text{cum}}\ G^{G}_{f}\ f^{-1} \tag{S1b}$$

$$\widehat{\mu} = \dot{\widehat{X}}\ \widehat{X}^{-1} + \frac{f}{\widehat{V}} \tag{S1c}$$

$$\widehat{\pi} = \left(\dot{\widehat{P}} + \frac{f}{\widehat{V}}\ \widehat{P}\right)\ \widehat{X}^{-1} \tag{S1d}$$

$$\widehat{\gamma} = \left(\frac{f}{\widehat{V}}\left(G_{\text{f}} - \widehat{G}\right) - \dot{\widehat{G}}\right)\widehat{X}^{-1} \tag{S1e}$$

$$\widehat{\gamma^{\circ}} = \widehat{\gamma}\left(1 - \frac{\widehat{P}}{\widehat{X}}\right)^{-1} \tag{S1f}$$

$$\widehat{\gamma^{\pi}} = \frac{\widehat{\pi}}{Y_{\frac{P}{G}}}\left(1 - \frac{\widehat{P}}{\widehat{X}}\right)^{-1} \tag{S1g}$$

$$\widehat{\gamma^{\mu}} = \left(\widehat{\mu} - \widehat{\pi}\ Y_{\frac{P}{G}}\right) Y_{\frac{X_{\text{r}}}{G}}{}^{-1}\left(1 - \frac{\widehat{P}}{\widehat{X}}\right)^{-1} \tag{S1h}$$

$$\widehat{\gamma^{\alpha}} = \widehat{\gamma^{\circ}} - \widehat{\gamma^{\mu}} - \widehat{\gamma^{\pi}} \tag{S1i}$$

$\widehat{\gamma^{\circ}}$ can only be estimated if the substrate concentration is above the limit of quantification.

To negate the effect of sampling we transform the feed and the volume by the sampling factor $s$. This creates an equivalent process without sampling where all concentrations $(\widehat{X}, \widehat{P}, \widehat{G})$ and the specific feed $(f/\widehat{V})$ are equal.

$$s(t) = \prod_{t_i \in S, t_i < t}\left(1 - \frac{V_{\text{sample}}}{\widehat{V}(t_i)}\right) \tag{S2a}$$

$$V_{\text{corr}}(t) = \frac{\widehat{V}(t)}{s(t)} \tag{S2b}$$

$$f_{\text{corr}}(t) = \frac{\widehat{f}(t)}{s(t)} \tag{S2c}$$

$S$ is the set of all sample times and $V_{\text{sample}}$ the sampling volume. Fitting and optimization are done for this corrected, unsampled process, and the results are transformed back to the sampled process.

#### S1.2 Cross validation and hyperparameter selection

In Section 2.1.2 the significance level $\alpha$ is treated as a hyperparameter and has to be defined by cross-validation, e.g., it depends on the sampling regime, more sampling points will require a lower $\alpha$. For cross-validation, the leave-one-out strategy estimates each run in the training data with the 11 other training runs. For these prediction, $R^2$ is calculated according to (12).

We repeat the model fit and cross-validation for different levels of significance (0.0001, 0,0002, 0.0005, 0,001, 0,002, 0,005, 0.01, 0.02, 0.05, 0.1, 0.2, 0.3, 0.4) and select the $\alpha$ with the highest $R^2$ for $P/X$ to determine the simplified model. For the case study an $\alpha$ of 0.02 is selected (Supplementary Figure S2).

### S1.3 Validation Simulations

In addition to the experimental validation, we add additional validation with simulated data.

#### *S1.3.1 Simulate experiments*

For the validation simulations, we generated 400 randomly reduced model equations (initial models). Although this meant that some parameters of the general model were randomly missing, certain parameters are always present as they comprise the simplest viable process model:

- $\gamma^{\circ}_{\max}$ and $K^{\circ}_{m}$ to describe the uptake solely dependent on the substrate concentration without inhibition,
- $\gamma^{\alpha}_{\min}$ and $K^{\alpha}_{g}$ to describe a linear maintenance function dependent on the uptake rate, and
- $\gamma^{\pi}_{\max}$ and $K^{\pi}_{m}$ to describe the production, dependent on the available substrate flux without inhibition.

In addition to these 6 necessary parameters, the model can include up to 15 additional terms, 4 inhibiting uptake, 4 inhibiting production, 4 increasing maintenance, and 3 adding temperature dependence. Each temperature dependence adds three additional model parameters while other terms add one model parameter (see Table S1). For each possible number of additional terms ($i = 0$ to 15), we randomly select 25 sets of parameters (choosing $i$ terms from the 15 available terms) and generate random values for these parameters. To get a reasonable range for inhibition and activation, the parameter value distribution was defined such that its median halves the product or uptake rate and doubles the maintenance requirement (detailed ranges are listed in Supplementary Table S5).

Subsequently, for all these initial models values for $X$, $P$, $G$, and $V$ are calculated for all runs and all measurement points according to the DoE, used for the training data, with the model equations (1-4). The initial biomass was set to 30 $\mathrm{g\,L^{-1}}$, i.e., the mean of the center point experiments. Parameterized models that do not reach a product concentration of at least 3 $\mathrm{g\,L^{-1}}$ in any process of the DoE are discarded and replaced by new equations created in the same way.

Next, we added randomly sampled errors to the biomass and product concentrations, mimicking measurement data with biological variance, sampling, and measurement errors.

$$\widehat{X} = X(1 + \epsilon_X), \quad \epsilon_X \sim \mathcal{N}(0, \sigma_X), \quad \widehat{P} = P(1 + \epsilon_P) \quad \epsilon_P \sim \mathcal{N}(0, \sigma_P) \tag{S3}$$

where $\mathcal{N}(0, \sigma)$ is the normal distribution and $\sigma_X \in \{0, 0.015, 0.03, 0.06\}$ and $\sigma_P \in \{0, 0.02, 0.04, 0.08, 0.16, 0.32\}$. 0.03 and 0.08 are the observed errors in the training data (see S1.3.3). Other values are added to explore the effect of errors on the OptFed results.

#### *S1.3.2 Compare OptFed results with initial models*

For all simulated experimental datasets we use OptFed to fit the model, including selecting relevant parameters. For each initial model, we calculate the optimal feed rate and temperature with OptFed and with RSM (exponential feeds are used to reduce the computational effort and make OptFed results comparable to RSM results). This gives us three sets of control variables ($\mu_f$ and $T$), and for each one, we calculate the final product-to-biomass yield using the initial model. The ratio between the optimum achieved by using the model prediction and the real optimum for the initial model is used as the performance indicator for the RSM and OptFed.

$$r_{\text{opt}}^{\text{OptFed}} = \frac{\mathcal{Y}_{\frac{P}{X}}\left(\mu_{f,\text{opt}}^{\text{OptFed}}, T_{\text{opt}}^{\text{OptFed}}\right)}{\mathcal{Y}_{\frac{P}{X}}\left(\mu_{f,\text{opt}}, T_{\text{opt}}\right)} \qquad r_{\text{opt}}^{\text{RSM}} = \frac{\mathcal{Y}_{\frac{P}{X}}\left(\mu_{f,\text{opt}}^{\text{RSM}}, T_{\text{opt}}^{\text{RSM}}\right)}{\mathcal{Y}_{\frac{P}{X}}\left(\mu_{f,\text{opt}}, T_{\text{opt}}\right)} \tag{S4}$$

A ratio close to 1 means we get very close to the optimum regarding the target variable $P/X$.

#### *S1.3.3 Estimating Variance in the training data*

We require sensible estimates of the expected errors to simulate meaningful experimental data. This is done using the four center point runs of the training data. These runs have identical uptake rates (feed rate is adapted to biomass concentration and volume) but different initial biomass concentrations and volumes. We compensate for this by calculating adjusted product and biomass concentrations for the process $i$ ($X_{\text{adj}}^i$ and $P_{\text{adj}}^i$, $i \in \mathcal{P}$) independent of the initial biomass concentration ($X_0$) and the initial volume ($V_0$). For $x \in \{X, P\}$ we caclulate

$$x_{\text{adj}}^i(t) = \frac{x^i(t)V^i(t)}{X_0^i V_0^i} \tag{S5a}$$

and based on this the relative deviation from the mean at each time-point

$$x_{\text{err}}^i(t) = \frac{x_{\text{adj}}(t) - \frac{1}{|\mathcal{P}|}\sum_{i\in\mathcal{P}} x_{\text{adj}}^i(t)}{\frac{1}{|\mathcal{P}|}\sum_{i\in\mathcal{P}} x_{\text{adj}}^i(t)} \tag{S5b}$$

for each process the mean error and the variance are calculated over all measurements points $\mathcal{M}$

$$\overline{x_{\text{err}}^i} = \frac{1}{|\mathcal{M}|}\sum_{t_j\in\mathcal{M}} x_{\text{err}}^i(t_j) \tag{S5c}$$

$$s^2(x_{\text{err}}^i) = \frac{1}{|\mathcal{M}|-1}\sum_{t_j\in\mathcal{M}} (X_{\text{err}}^i(t_j))^2 \tag{S5d}$$

and use the mean of the variance to characterize the random error expected within a process.

$$s^2(x_{\text{err}}) = \frac{1}{|\mathcal{P}|} \sum_{i \in \mathcal{P}} s^2(X^i_{\text{err}}) \tag{S5e}$$

The expected random error within one process in (S5e) is critical to fit the uptake, growth, and production rates and the model equations. We use this to assess the simulated validation results.

#### S1.4 Temperature stability

To estimate the stability of our temperature optimum we use a strategy based on the Monte Carlo method [64–66], using the estimated relative errors of our data determined in Section S1.3.3 (3 % for biomass and 8 % for product). We create 1000 disturbed datasets according to Eq. (S3) and fit each set's model for $\gamma^\pi$. For each model, we calculate the optimal temperature and intervals where $p$ % of the maximum is reached ($0 < p < 100$). We now search for the maximal ratio and the corresponding temperature we can guarantee statistically significant ($\alpha = 0.05$). To do this, we determine the temperature for each ratio $p$ where the least number of models is outside the temperature ratio. We search for the highest ratio ($\gamma^\pi(T)/\gamma^\pi(t_{\text{opt}})$) where more than 95 % of all processes reach this ratio at the optimal temperature. This temperature is the stable optimum.

### S2 Supplementary Tables

Supplementary Table S1: List of parameters used throughout the manuscript. Values shows the parameter values identified by OptFed. Where no value is given, this parameter was removed during model simplification. Increased error gives the additional fitting error (calculated as RSS, see (12)) if this parameter would be removed as well.

| | Name | Unit | Value | increased error |
|---|---|---|---|---|
| Parameters (fitted using training data) | | | | |
| *Parameters of base model* | | | | |
| $c_{\gamma^{\circ}_{\text{max}}}$ | maximal uptake rate | $\text{g}\,\text{g}^{-1}\,\text{h}^{-1}$ | 0.49 | |
| $K^{\circ}_{m}$ | dependence on substrate concentration | g | $1.0 \times 10^{-3}$ | |
| $c_{\gamma^{\alpha}_{\text{min}}}$ | maintenance requirement without growth | $\text{g}\,\text{g}^{-1}\,\text{h}^{-1}$ | $2.4 \times 10^{-5}$ | |
| $K^{\alpha}_{g}$ | growth dependent maintenance | $\text{g}\,\text{g}^{-1}\,\text{h}^{-1}$ | $1.0 \times 10^{-4}$ | 142 % |
| $c_{\gamma^{\pi}_{\text{max}}}$ | maximal production rate | $\text{g}\,\text{g}^{-1}\,\text{h}^{-1}$ | - | |
| $K^{\pi}_{m}$ | production dependence on available substrate | $\text{g}\,\text{g}^{-1}\,\text{h}^{-1}$ | 0.62 | 188 % |
| inhibition parameter for $\gamma^{\circ}$ | | | | |
| $K^{\circ}_{G}$ | substrate inhibition | $\text{g}\,\text{L}^{-1}$ | 89 | 119 % |
| $K^{\circ}_{n}$ | inhibition through no. of generations | | - | |
| $K^{\circ}_{P}$ | product inhibition | $\text{g}\,\text{L}^{-1}$ | - | |
| $K^{\circ}_{X}$ | biomass inhibition | $\text{g}\,\text{g}^{-1}$ | - | |
| parameters for increased $\gamma^{\alpha}$ | | | | |
| $K^{\alpha}_{G}$ | increase caused by free substrate | $\text{g}\,\text{L}^{-1}$ | - | |
| $K^{\alpha}_{n}$ | increase with increasing no. of generations | | - | |
| $K^{\alpha}_{P}$ | increase caused by product | $\text{g}\,\text{L}^{-1}$ | 0.11 | 37 % |
| $K^{\alpha}_{X}$ | increase caused by biomass | $\text{g}\,\text{g}^{-1}$ | - | |
| inhibition parameters for $\gamma^{\pi}$ | | | | |
| $K^{\pi}_{G}$ | effect of available substrate on production | $\text{g}\,\text{L}^{-1}$ | - | |
| $K^{\pi}_{n}$ | inhibition for higher no. of generations | | 1.5 | 50 % |
| $K^{\pi}_{P}$ | Product inhibition | $\text{g}\,\text{g}^{-1}$ | - | |
| $K^{\pi}_{X}$ | Biomass inhibition | $\text{g}\,\text{L}^{-1}$ | - | |
| *Temperature dependence for* $\gamma^{\circ}$ | | | | |
| $\Delta G^{\circ}_{\text{cat}}$ | catalytic activation energy | $\text{J}\,\text{mol}^{-1}$ | - | |
| $\Delta H^{\circ}_{\text{eq}}$ | enthalpic (conversion of active to inactive enzyme) | $\text{J}\,\text{mol}^{-1}$ | - | |
| $T^{\circ}_{\text{eq}}$ | temperature where half the enzyme is active | K | - | |
| $E^{\circ}_{0}$ | hypothetical enzyme concentration | K | - | |
| *Temperature dependence for* $\gamma^{\alpha}$ | | | | |
| $\Delta G^{\alpha}_{\text{cat}}$ | catalytic activation energy | $\text{J}\,\text{mol}^{-1}$ | - | |
| $\Delta H^{\alpha}_{\text{eq}}$ | enthalpic (conversion of active to inactive enzyme) | $\text{J}\,\text{mol}^{-1}$ | - | |
| $T^{\alpha}_{\text{eq}}$ | temperature where half the enzyme is active | K | - | |
| $E^{\alpha}_{0}$ | hypothetical enzyme concentration | K | - | |
| *Temperature dependence for* $\gamma^{\pi}$ | | | | |
| $\Delta G^{\pi}_{\text{cat}}$ | catalytic activation energy | $\text{J}\,\text{mol}^{-1}$ | $5.2 \times 10^{4}$ | 38 % |
| $\Delta H^{\pi}_{\text{eq}}$ | enthalpic (conversion of active to inactive enzyme) | $\text{J}\,\text{mol}^{-1}$ | $4.8 \times 10^{6}$ | |
| $T^{\pi}_{\text{eq}}$ | temperature where half the enzyme is active | K | 310 | |
| $E^{\pi}_{0}$ | hypothetical enzyme concentration | K | $9.8 \times 10^{-9}$ | |
| *Constants* (physical constants and yield determined by metabolomic model | | | | |
| $Y_{\frac{X_r}{G}}$ | biomass yield per substrate | $\text{g}\,\text{g}^{-1}$ | 0.627 | |
| $Y_{\frac{P}{G}}$ | product yield per substrate | $\text{g}\,\text{g}^{-1}$ | 0.652 | |
| $k_B$ | Boltzmann constant | $\text{J}\,\text{K}^{-1}$ | $1.38 \times 10^{-23}$ | |
| $R$ | Gas constant | $\text{J}\,\text{mol}^{-1}\,\text{K}^{-1}$ | 8.314 | |
| $h$ | Plank constant | $\text{J}\,\text{h}$ | $2.39 \times 10^{-30}$ | |

Supplementary Table S2: Maximal and minimal values for all fitted parameters.

| Parameter | min | max |
|---|---|---|
| $K_m^\circ$ | $1 \times 10^{-15}$ | $1 \times 10^{2}$ |
| $K_X^\circ$ | 0 | $1 \times 10^{4}$ |
| $K_P^\circ$ | 0 | $1 \times 10^{4}$ |
| $K_n^\circ$ | 0 | $1 \times 10^{4}$ |
| $K_G^\circ$ | $1 \times 10^{-5}$ | $1 \times 10^{4}$ |
| $c_{\gamma^\circ_{\text{max}}}$ | 0 | 1 |
| $E_0^\circ$ | 0 | $1 \times 10^{-5}$ |
| $\Delta G_{\text{cat}}^\circ$ | $1 \times 10^{4}$ | $1 \times 10^{5}$ |
| $\Delta H_{\text{eq}}^\circ$ | $1 \times 10^{5}$ | $1 \times 10^{7}$ |
| $T_{\text{eq}}^\circ$ | 300 | 315 |
| $K_g^\alpha$ | $1 \times 10^{-6}$ | $\infty$ |
| $K_X^\alpha$ | $1 \times 10^{-6}$ | $\infty$ |
| $K_P^\alpha$ | $1 \times 10^{-6}$ | $\infty$ |
| $K_n^\alpha$ | $1 \times 10^{-6}$ | $\infty$ |
| $K_G^\alpha$ | $1 \times 10^{-6}$ | $\infty$ |
| $c_{\gamma^\alpha_{\text{min}}}$ | 0 | 1 |
| $E_0^\alpha$ | 0 | $1 \times 10^{-5}$ |
| $\Delta G_{\text{cat}}^\alpha$ | $1 \times 10^{4}$ | $1 \times 10^{5}$ |
| $\Delta H_{\text{eq}}^\alpha$ | $1 \times 10^{5}$ | $1 \times 10^{7}$ |
| $T_{\text{eq}}^\alpha$ | 300 | 315 |
| $K_m^\pi$ | 0 | $1 \times 10^{2}$ |
| $K_X^\pi$ | 0 | $1 \times 10^{4}$ |
| $K_P^\pi$ | 0 | $1 \times 10^{4}$ |
| $K_n^\pi$ | 0 | $1 \times 10^{4}$ |
| $K_G^\pi$ | 0 | $1 \times 10^{4}$ |
| $c_{\gamma^\pi_{\text{max}}}$ | 0 | 1 |
| $E_0^\pi$ | 0 | $1 \times 10^{-5}$ |
| $\Delta G_{\text{cat}}^\pi$ | $1 \times 10^{4}$ | $1 \times 10^{5}$ |
| $\Delta H_{\text{eq}}^\pi$ | $1 \times 10^{5}$ | $1 \times 10^{7}$ |
| $T_{\text{eq}}^\pi$ | 300 | 315 |

Supplementary Table S3: Estimated parameters for the response surface methodology (RSM) model according to Eqn. (11).

| Parameter | $c$ | $c_{\text{f}}$ | $c_{\text{T}}$ | $c_{\text{fT}}$ | $c_{\text{f}^2}$ | $c_{\text{T}^2}$ |
|---|---|---|---|---|---|---|
| Value | $-2.11 \times 10^{2}$ | $-16.7$ | 1.39 | 0.0607 | $-7.45$ | $-2.30 \times 10^{-3}$ |
| p-Value | 0.20 | 0.54 | 0.20 | 0.50 | 0.055 | 0.20 |

Supplementary Table S4: Performance summary of all processes (identified by ID) designed and analyzed in this work. For 3 temperatures, 31.0 °C (center point of training data), 33.5 °C, and 35.8 °C (optimal temperature) the optimum was calculated with linear feed function (opt. process) and free feed function.

| ID | comment | feed rate, $f$ [$L\,h^{-1}$] | $P$ [$g\,L^{-1}$] | $\Delta P$ [%] | $P/X$ [$g\,g^{-1}$] | $\Delta P/X$ [%] | $X_{end}$ [$g\,L^{-1}$] | $V/V_0$ |
|---|---|---|---|---|---|---|---|---|
| 0 | reference | - | 7.76 | 0.00 | 0.14 | 0.00 | 57.11 | 1.42 |
| 1 | measurements | - | 7.67 | −0.01 | 0.13 | −0.04 | 58.47 | 1.29 |
| | $T = 31.0\,°C$ | | | | | | | |
| 2 | opt process | $0.02208 + 0.00377t$ | 7.51 | −3.24 | 0.14 | 0.34 | 55.07 | 1.35 |
| 5 | free feed | - | 7.51 | −3.23 | 0.14 | 0.34 | 55.07 | 1.35 |
| | $T = 33.5\,°C$ | | | | | | | |
| 3 | opt process | $0.02190 + 0.00443t$ | 8.65 | 11.47 | 0.16 | 18.99 | 53.50 | 1.38 |
| 6 | free feed | - | 8.65 | 11.48 | 0.16 | 18.99 | 53.50 | 1.38 |
| | $T = 35.8\,°C$ | | | | | | | |
| 4 | opt process | $0.02154 + 0.00526t$ | 9.64 | 24.23 | 0.19 | 37.08 | 51.76 | 1.42 |
| 7 | free feed | - | 9.64 | 24.22 | 0.19 | 37.08 | 51.75 | 1.42 |

Supplementary Table S5: Distribution of random parameter values for simulated models. The range of basic parameter values is based on the values observed in the training data. Typically an active parameter should halve the uptake or production rate for the highest observed values or double the maintenance. The temperature range is selected to reflect similar behavior to the reference processes with an optimum within the design space of the DoE.

| Basic parameters: | | |
|---|---|---|
| $c_{\gamma^{\circ}_{\text{max}}}$ | uniformly distributed | $[0.3,\ 1]$ g g$^{-1}$ h$^{-1}$ |
| $K^{\circ}_{m}$ | uniformly distributed | $[0,\ 1]$ g |
| $c_{\gamma^{\alpha}_{\text{min}}}$ | uniformly distributed | $[0,\ 0.05]$ g g$^{-1}$ h$^{-1}$ |
| $K^{\alpha}_{g}$ | uniformly distributed | $[0,\ \frac{1}{2\gamma^{\alpha}_{\text{min}}}]$ $(\text{g g}^{-1}\ \text{h}^{-1})^{-1}$ (0 to 50 % of total uptake) |
| $c_{\gamma^{\pi}_{\text{max}}}$ | uniformly distributed | $[0,\ 0.1]$ g g$^{-1}$ h$^{-1}$ |
| $K^{\alpha}_{g}$ | exponential distributed | mean $= 1$ g g$^{-1}$ h$^{-1}$ |
| Temperature (for $i \in \{\mu, \pi, \circ\}$m $j \in \{\max, \min\}$) : | | |
| $\Delta G^{i}_{\text{cat}}$ | uniformly distributed | $[4 \times 10^4,\ 1 \times 10^5]$ mol J$^{-1}$ |
| $\Delta H^{i}_{\text{eq}}$ | uniformly distributed | $[7 \times 10^4,\ 1 \times 10^6]$ mol J$^{-1}$ |
| $T^{i}_{\text{eq}}$ | uniformly distributed | $[304.15,\ 308.15]$ K |
| $E^{i}_{0}$ | calculate from $c_{\gamma^{i}_{j}}$ | $c_{\gamma^{i}_{j}} = \gamma^{i}_{j}(304.15)$ K |
| inhibition parameters for $g$ | | |
| $K^{\circ}_{G}$ | exponentially distributed | median $= \max(G)$ [g L$^{-1}$] |
| $K^{\circ}_{n}$ | exponentially distributed | median $= \max(n)$ |
| $K^{\circ}_{P}$ | exponentially distributed | median $= \max(P)$ [g g$^{-1}$] |
| $K^{\circ}_{X}$ | exponentially distributed | median $= \max(X)$ [g L$^{-1}$] |
| parameters for increased $\gamma^{\alpha}$ | | |
| $K^{\alpha}_{G}$ | exponentially distributed | median $= (\max(G))^{-1}$ [g L$^{-1}$] |
| $K^{\alpha}_{n}$ | exponentially distributed | median $= \max(n)$ |
| $K^{\alpha}_{P}$ | exponentially distributed | median $= \max(P)$ [g g$^{-1}$] |
| $K^{\alpha}_{X}$ | exponentially distributed | median $= \max(X)$ [g L$^{-1}$] |
| inhibition parameters for $\gamma^{\pi}$ | | |
| $K^{\pi}_{G}$ | exponentially distributed | median $= \max(G)$ [g L$^{-1}$] |
| $K^{\pi}_{n}$ | exponentially distributed | median $= \max(n)$ |
| $K^{\pi}_{P}$ | exponentially distributed | median $= \max(P)$ [g g$^{-1}$] |
| $K^{\pi}_{X}$ | exponentially distributed | median $= \max(X)$ [g L$^{-1}$] |

### S3 Supplementary Figures

**Author details**
[1]Department of Analytical Chemistry, University Vienna, Währinger Straße, 1090 Vienna, Austria, EU. [2]Doctorate School of Chemistry, University of Vienna, Währinger Straße, 1090 Vienna, Austria, EU. [3]Integrated Bioprocess Development, Technical University Vienna, Getreidemarkt 9, 1060 Vienna, Austria, EU. [4]Austrian Centre of Industrial Biotechnology, Krenngasse 37, 8010 Graz, Austria, EU.

**References**

1. Stefan Kittler, Julian Ebner, Mihail Besleaga, Johan Larsbrink, Barbara Darnhofer, Ruth Birner-Gruenberger, Silvia Schobesberger, Christopher K Akhgar, Andreas Schwaighofer, Bernhard Lendl, et al. Recombinant protein I: production, purification and characterization of a universal binding ligand. *Journal of Biotechnology*, 359: 108–115, 2022.

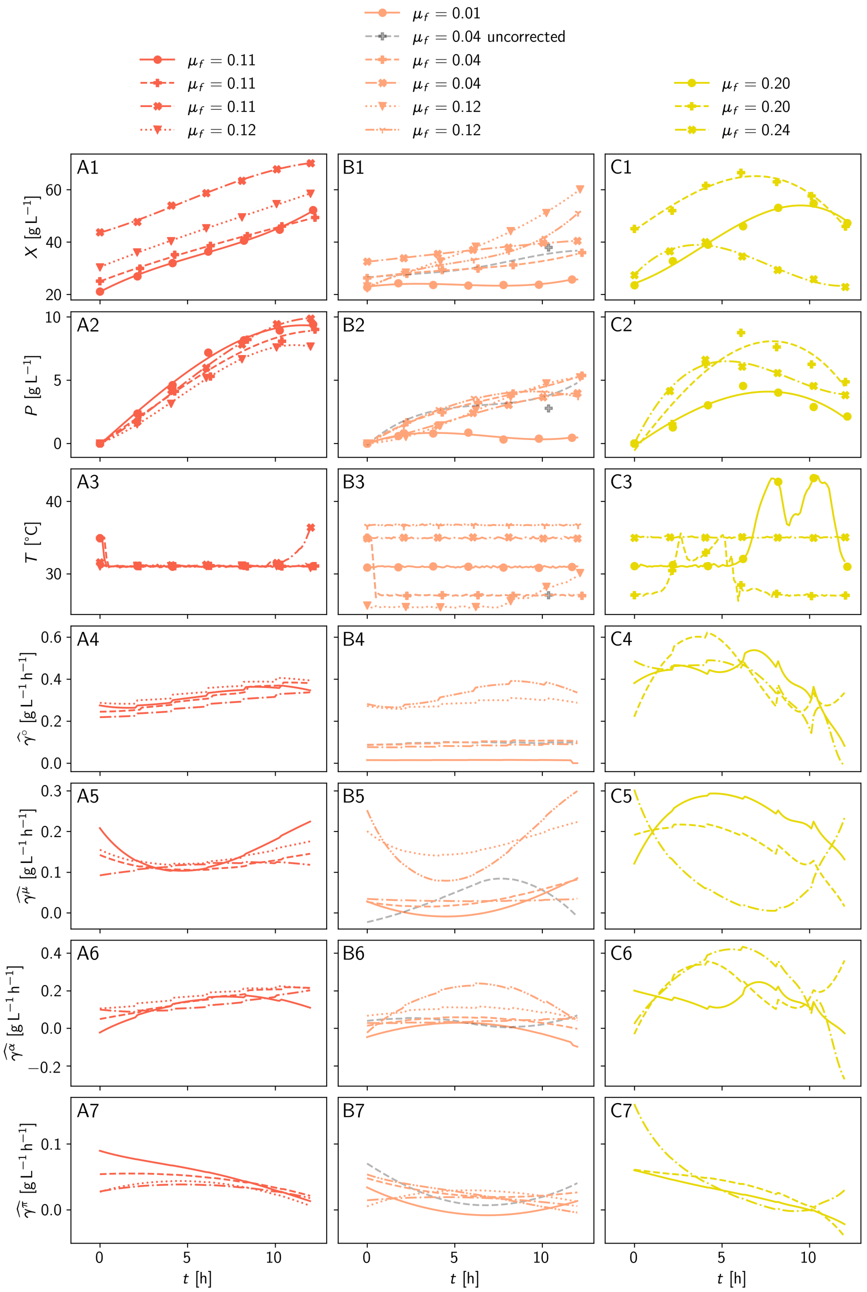


Supplementary Figure S1: **Fitted Splines.** The fitted splines (lines) generate stable rates from noisy experimental data (markers). Biomass (row 1) and product (row 2) concentrations allow the calculation of stable uptake rates (row 4) and substrate usage rates (rows 5 - 7). Column (A) shows the repeats at the center points, (B) other processes without substrate accumulation, and (C) processes with substrate accumulation. In column (B) one measurement at 10 hours is out of trend. This measurement was removed from the dataset. The fit including this point is shown in gray.

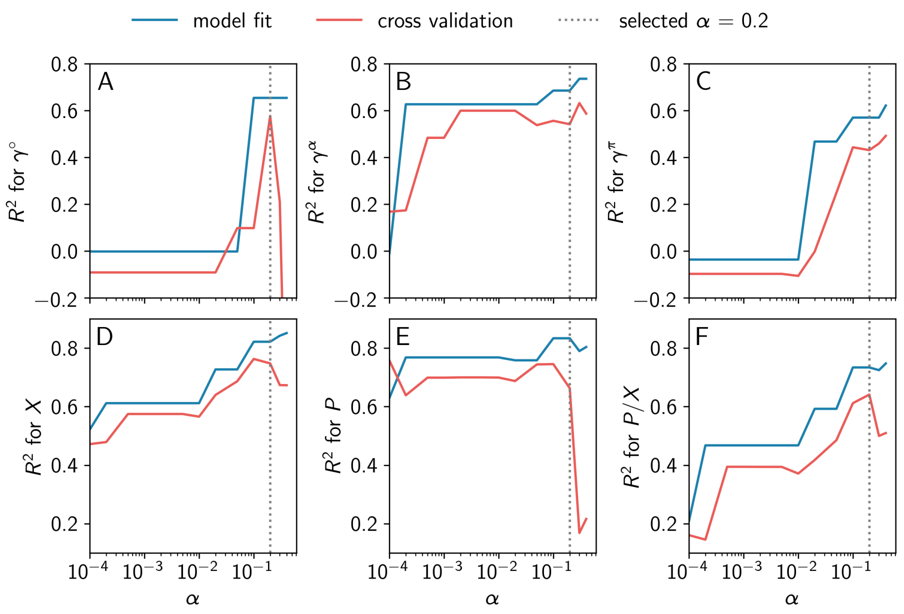


Supplementary Figure S2: **Hyperparameter selection.** The complexity of the used error depends on the significance level ($\alpha$) used in the algorithm described in Section 2.1.2. Here we show $R^2$, dependent on the significance level, for the fitted uptake rate (A) and substrate-to-maintenance flux (B) and the substrate-to-product flux (C), as well as the measured biomass and product concentrations (D and E) and product per biomass ratio (F). While $R^2$ for the model fit increases with model complexity (blue), it decreases in cross-validation (red). To get the best possible fit and avoid overfitting we choose a significance level of $\alpha = 0.2$. At this point, the highest $R^2$ for our target variable ($P/X$) is reached in cross-validation.

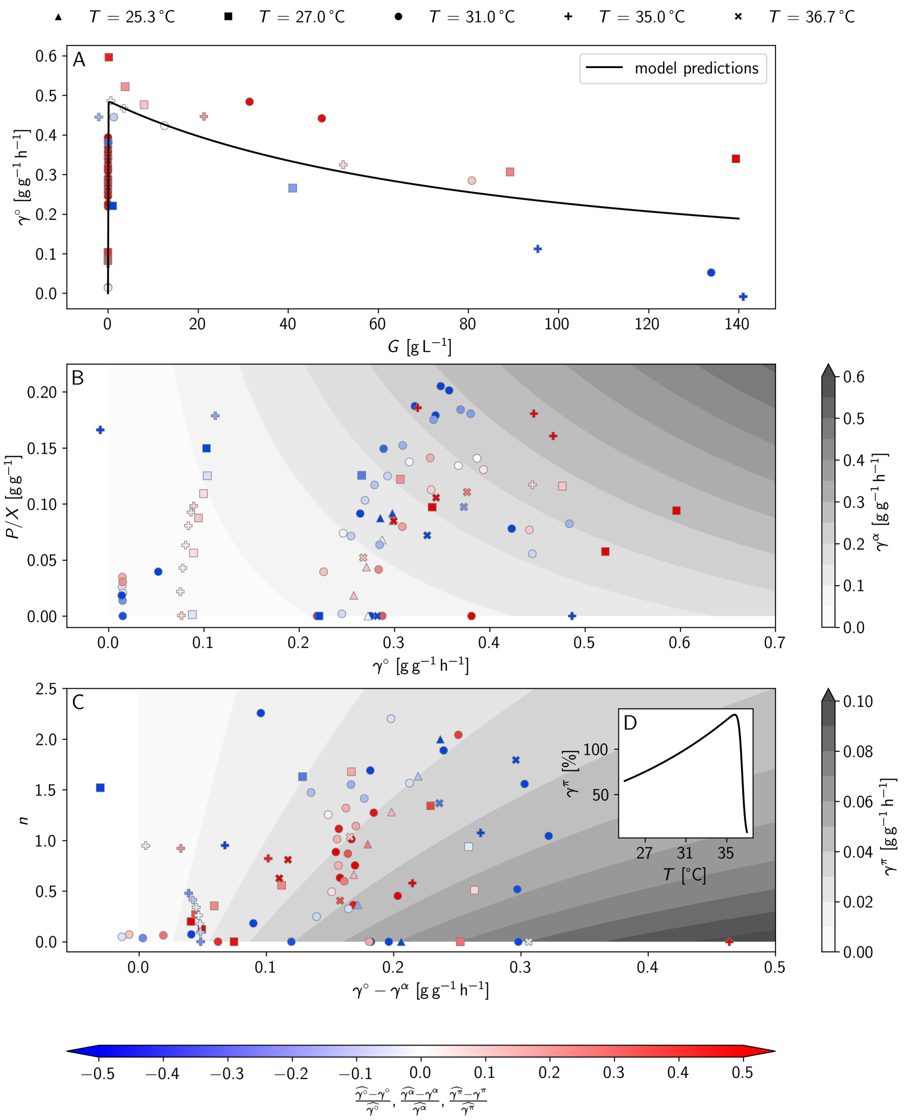


Supplementary Figure S3: **Model estimations** for the substrate uptake rate ($\gamma^\circ$, panel A, y-axis), the substrate-to-maintenance flux ($\gamma^\alpha$, panel B, shades of grey), and the substrate-to-product flux ($\gamma^\pi$, panel C, shades of grey). Marker shapes indicate the process' target temperature. Marker colors indicate model error as calculated by Eq. 13. $\gamma^\circ$ (A) solely depends on the substrate concentration. The maximum uptake is reached for low substrate concentrations and is inhibited at higher concentrations. $\gamma^\alpha$ (B) depends on the $\gamma^\circ$ and the product per biomass ratio. Both increase the required maintenance. $\gamma^\pi$ (C) depends on three variables, $\gamma^\circ - \gamma^\alpha$, $n$, and $T$. $\gamma^\circ - \gamma^\alpha$ and the number of generations, $n$, are shown in (C), the relative effect of temperature, $T$, is depicted in (D). The values of $\gamma^\pi$ in (C) are shown for 31 °C. For other temperatures, they are multiplied by the relative effect shown in (D), meaning a reduction of 34 % at 25.3 °C and 25 % at 27.0 °C, an increase of 32 % at 35.0 °C, and a decrease of 53 % at 36.7 °C.

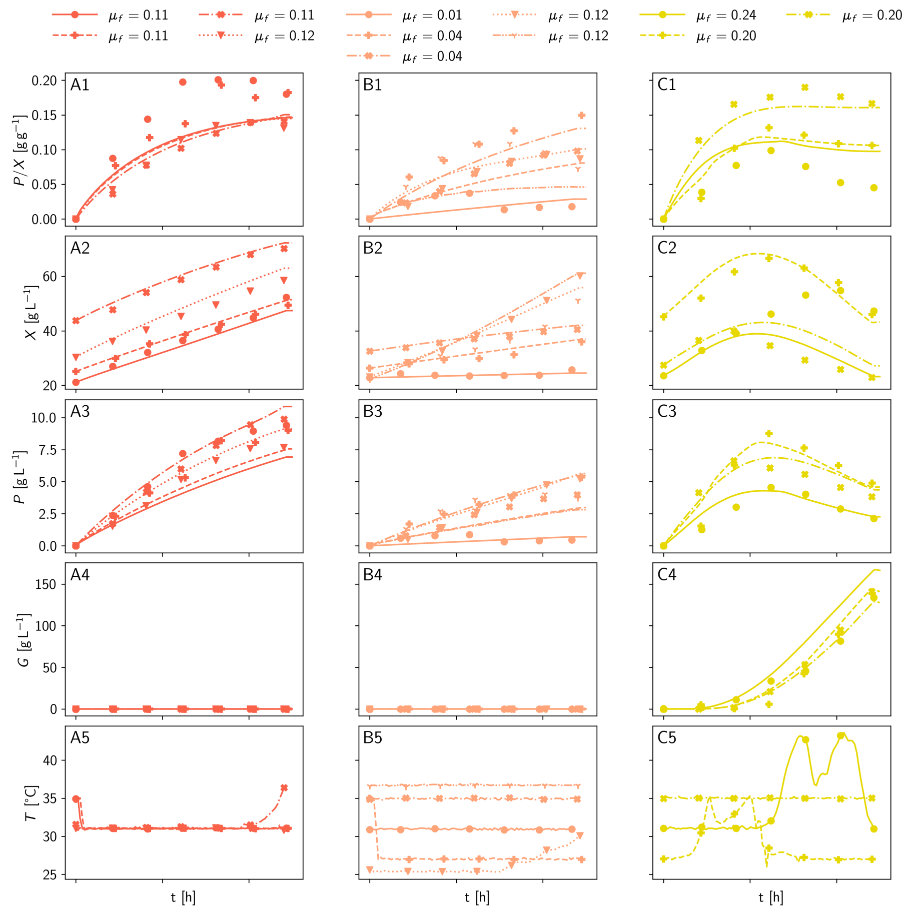


Supplementary Figure S4: **Reduced model predictions** (lines) are compared to measured data for product per biomass (row 1), biomass (row 2), product (row 3), substrate (row 4), and temperature (row 5). There is a good fit for processes without substrate accumulation (column A and B) and an adequate fit with substrate accumulation (C).

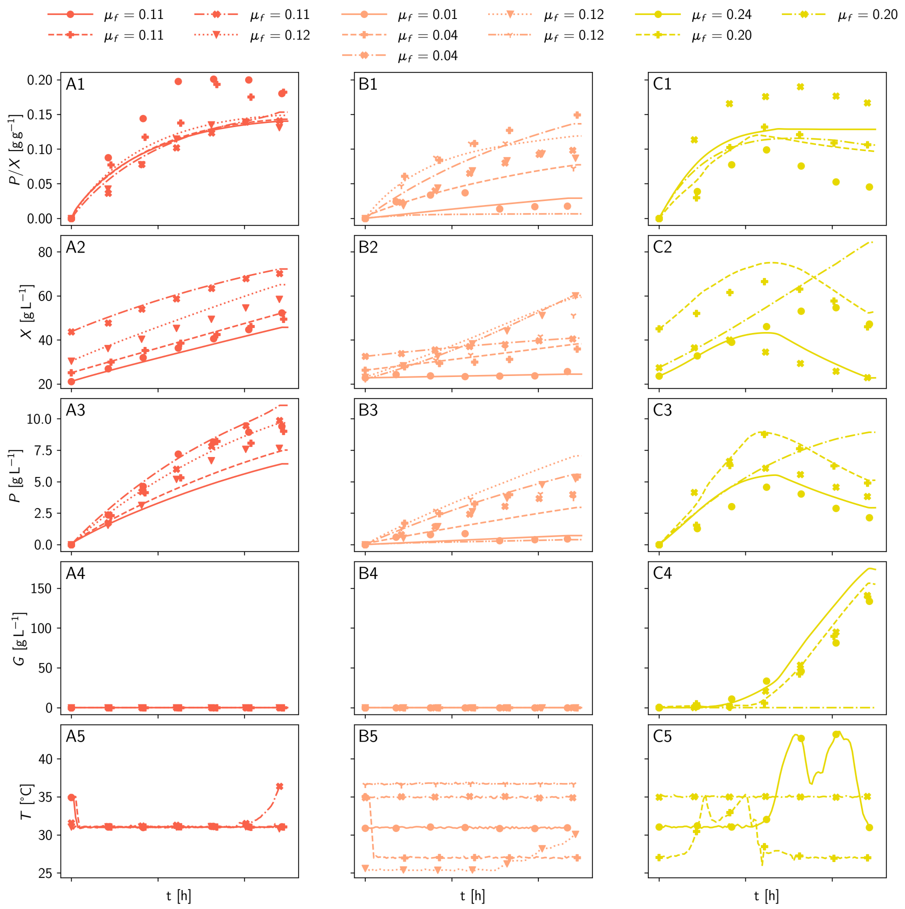


Supplementary Figure S5: **Cross-validation.** For cross-validation, the other 11 processes are used for model fitting, parameter reduction, and prediction (see S1.2). The fitting errors increase only moderately in cross-validation (see Table 2b and compare Supplementary Figure S4).

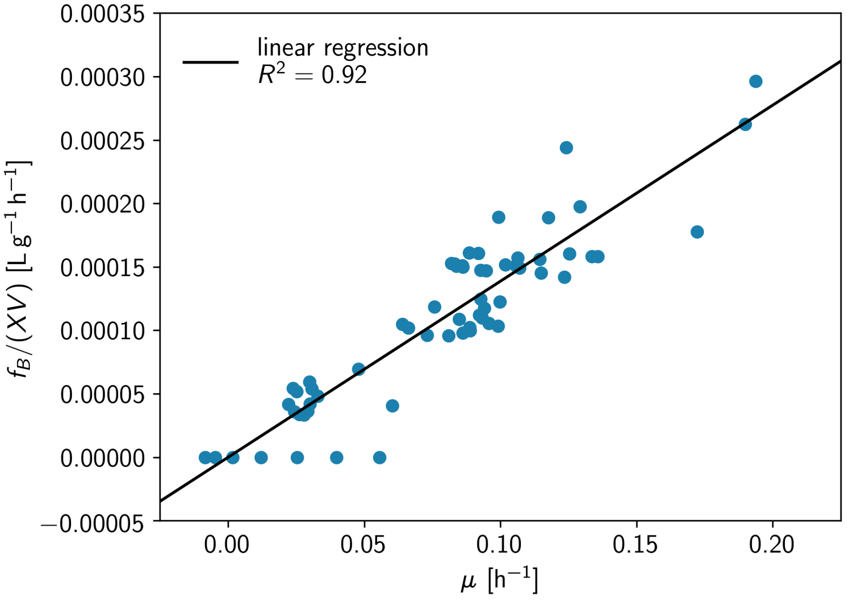


Supplementary Figure S6: **Base feed.** In addition to the substrate feed base (12.5 % $NH_4OH$) is added to control the pH value [1]. During optimization, the base requirement is estimated with a linear regression model based on the nine training processes without substrate accumulation.

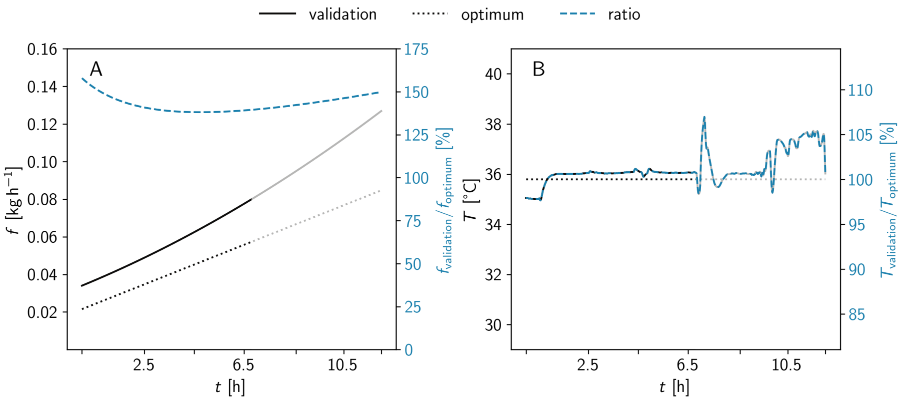


Supplementary Figure S7: **Comparison of the predicted (non-linear) and the experimental validation (linearized) process variables.** The feed rate ($f$) is shown in panel A, and the temperature ($T$) in panel B. The differences in the feed rate have a minor impact on the process performance. Figure 2 shows that the optimum is very broad for different feed rates. Even differences in the feed rate of 50 % have a minimal effect on the predicted product-to-biomass yield (less than 1 %). On the other hand, the temperature deviations are more detrimental, especially after 6.5 hours, they become a problem. Especially the temperature spikes, due to handling and an undersized cooling capacity may lead to biological changes that are not well reflected in our process model.

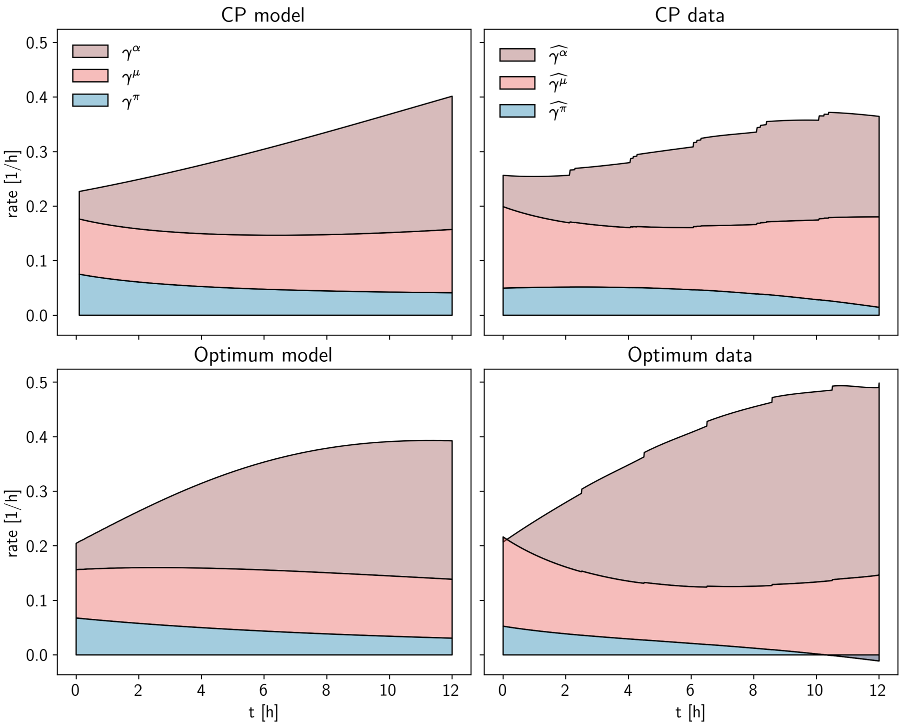


Supplementary Figure S8: **Flux distribution**. The substrate uptake ($\gamma^\circ$) is distributed between production (bottom), growth (center), and maintenance (top). This is shown for the center point of the training data (top row) and the predicted optimum (bottom row) as well as for the model prediction (left) and the experimental rates(right). Maintenance requirements with time, while production decreases. There is a good relation between the experimental data and the model predictions for the center points, where a mean of all four center points is used. For the optimum, the general trend still holds, but without repeats rate fits are less stable. The drop in production due to unstable conditions is clearly visible.

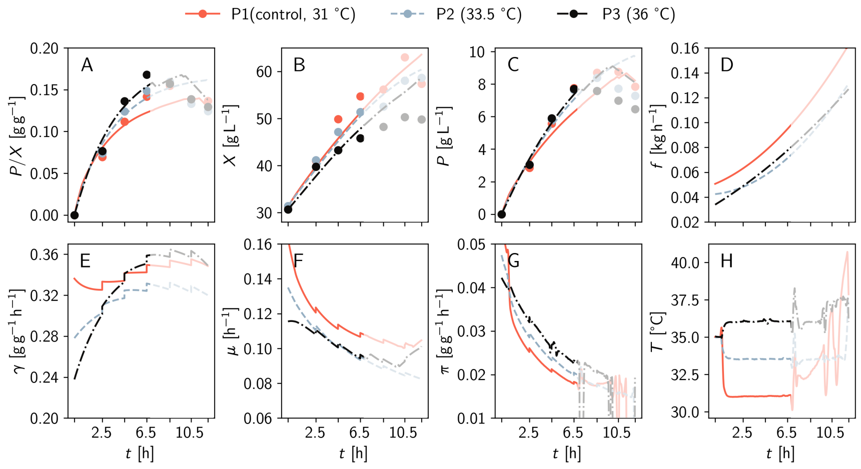


Supplementary Figure S9: **Validation of the predicted optimal fermentation.** In addition to the panels shown in Figure 4 (A-E, H), we show the feed (D), uptake (E), growth (F), and production (G) rate for the validation experiments.

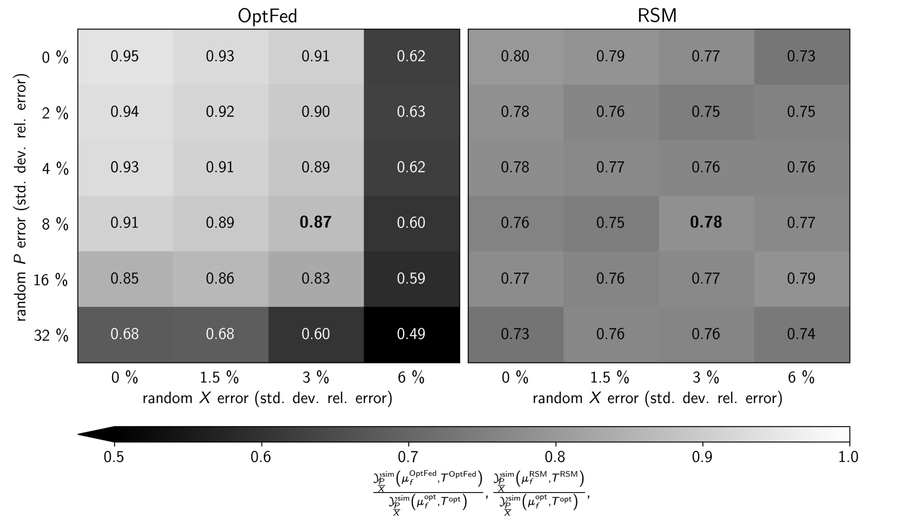


Supplementary Figure S10: **Computational validation.** Comparing the yields for all simulated processes ($\mathcal{Y}^{\text{sim}}_{\frac{P}{X}}$) at the optimum with the predicted optimum identified by OptFed. A ratio of 1 means that the optimum is correctly identified, and 0 means that no production happens at the predicted optimum. For process variation observed in the training data (bold, standard deviation of relative errors are 3 % for the biomass and 8 % for the product). RSM is very stable against variations in data, but can only reach a mean 78 % of the theoretical optimum compared to 87 % with OptFed at experimental error levels. The distribution of results is shown in Supplementary Figure S13. The creation and evaluation of the random processes are described in Section S1.3.

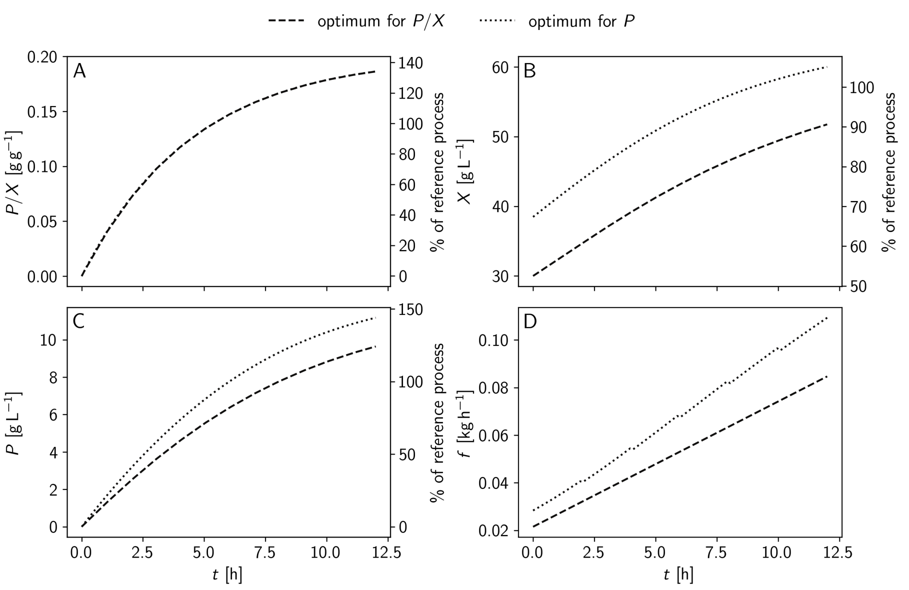


Supplementary Figure S11: **Comparison of $\mathcal{Y}_{\frac{P}{X}}$ and $PV$ optimized processes.** Given the process constraints and fitted model in our case study, the optimal processes are equal in $\mathcal{Y}_{\frac{P}{X}}$ and vary less than 16 % in $P$ of each other.

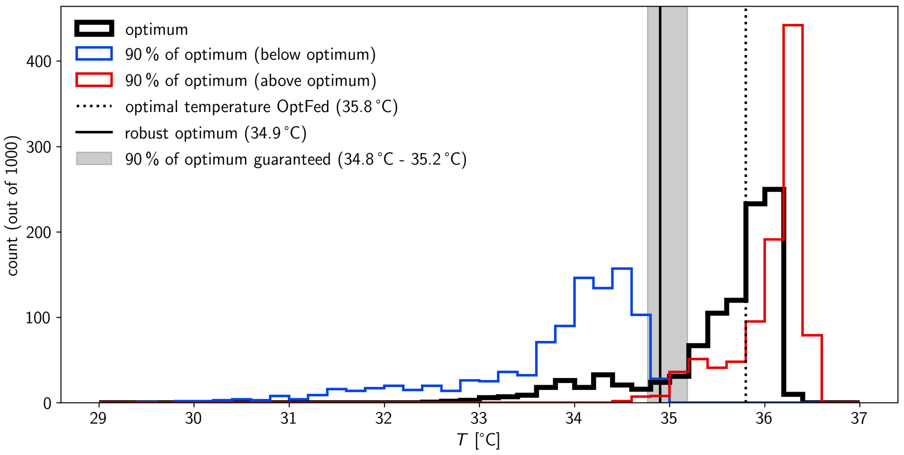


Supplementary Figure S12: To estimate the possible error of missing the optimal temperature we add errors according to Supplementary methods S1.3.3 and calculate the temperature dependence with these modified data points 1000 times. Reducing the temperature from 35.8 °C (dotted line) to 34.9 °C (solid line) reduced the production rate by 5 %, but guarantees that the models with disturbed data are all within 10 % of their optimum (at significance level 0.05). The black histogram shows the distribution of the optima while the blue and red histograms indicate the lower and upper bounds of the area with less than 10 % error. For temperatures in the gray array less than 5 % of the models reach less than 90 % of their optimum. Additional methodical information is found in Section S1.4.

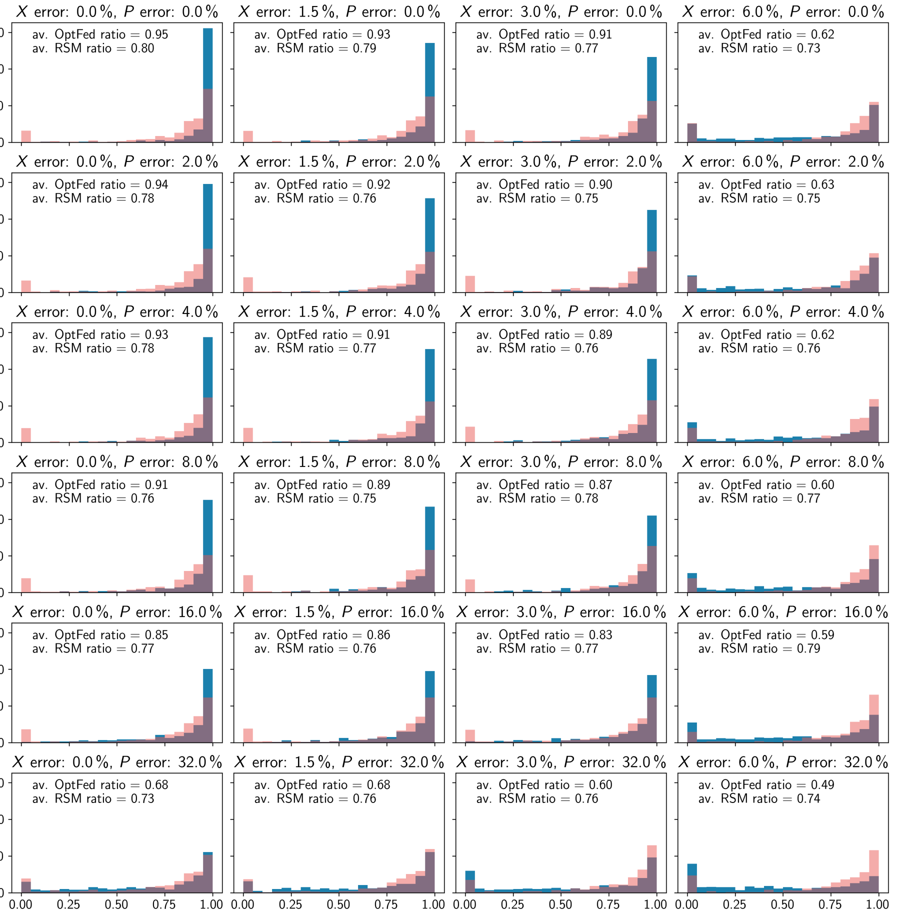


Supplementary Figure S13: **Computational validation / distributions.** In addition to the mean value of the optima ratio shown in Figure S10, we see the distribution of the ratios. For low and medium error levels OptFed the predicted optimum is very close to he model optimum and outliers are rare.
